# Root-associated microbial community recruitment in two citrus rootstocks subjected to water and salinity stresses

**DOI:** 10.64898/2026.08.06.743354

**Authors:** A. Mosca, G. Modica, G. Dimaria, D. Nicotra, M.F. Lombardo, G. Cirvilleri, A. Genitle, A. Pulvirenti, A. Continella, V. Catara

## Abstract

**Background and Aims:** Abiotic stress is a major constraint for citrus production in Mediterranean environments, where water deficit and salinity frequently occur. This is particularly relevant for perennial crops, like citrus, where limited options for stress avoidance exist. Rootstocks are extensively employed to enhance stress resilience; however, their influence on the root microbiome under abiotic stress remains largely unexplored. Here, we investigated the effects of water stress and salinity on the diversity, composition, and interactions of bacterial and fungal communities in two citrus rootstocks with reported contrasting phenotypes, such as Bitters, which has been described as exhibiting a promising tolerance to both water and salt stress, and Carrizo, which is generally reported to be highly sensitive to these conditions.

**Methods:** The distinct rootstocks have been subjected to either water stress or salt stress and compared with the non-stressed rootstocks. At the end of stress period, they were profiled and then integrated with recorded plant morphological (i.e. root volume), physiological (water potential, abscisic acid, chlorophyll and chlorophyll content meter) and biochemical measurements (abscisic acid and catalase). In parallel, we used a high-throughput amplicon sequencing to profile bacterial and fungal communities inhabiting the rhizosphere and endorhizosphere microhabitats of the rootstocks in both stresses and in non-treated conditions. Finally, we used correlations and multivariate analysis to determine relationships between plant performance and microbiome putatively underpinning stress adaptation and tolerance.

**Results:** Across all treatments, microbial community composition was primarily shaped by microhabitat, with clear differentiation between rhizosphere and endorhizosphere. Abiotic stress significantly restructured microbial communities, particularly in the rhizosphere, while the endorhizosphere exhibited stronger genotype-dependent patterns. Bacterial communities showed pronounced stress-driven enrichments of taxa belonging to the main phyla (such as Proteobacteria, Actinobacteriota and Bacteroidota), with selective recruitment of taxa putatively associated with stress adaptation, whereas the response of fungal taxa (more represented by Ascomycota, Basidiomycota and Glomeromycota phyla) was less consistent and mainly microhabitat-driven. Notably, the two rootstocks exhibited distinct physiological strategies, with Bitters by increased proline accumulation and root volume and Carrizo characterized by enhanced ABA and catalase.

**Conclusions:** Our findings showed Bitters outperform Carrizo in terms of tolerance to both water and salinity stress. In both rootstocks, specific bacterial taxa such as high abundant core or rare members, were associated with distinct phenotypic parameters, highlighting the importance of integrating plant and microbiome perspectives for improving stress resilience in citrus.

## Introduction

Citrus represents one of the most extensively cultivated fruit crops worldwide, with cultivation spanning over 140 countries. Within the European Union, total citrus production exceeded 10.6 million tons, with oranges alone accounting for more than 7.4 million tons. Italy ranks as the second largest producer of citrus in the EU, contributing over 3 million tons to the total production with oranges representing around 58% of the total Italian production, whereas lemon and limes around the 14% (FAOSTAT, 2024, accessed on June 14, 2026).

Citrus is predominantly cultivated in warm regions, including the Mediterranean basin, where limited water availability necessitates supplemental irrigation. However, irrigation practices often result in the accumulation of salts in the soil, leading to salinity stress (García-Sánchez et al., 2002). Citrus is widely recognized as a salt-sensitive crop (Maas 1993), and elevated soil salinity can severely impair plant growth, yield, and fruit quality as well as drought (Lo Piero 2020). These effects are mainly associated with osmotic and ionic stress, which disrupt cellular homeostasis and interfere with key physiological processes, including metabolic activity and hormonal regulation (García-Sánchez et al. 2002; Vanella et al. 2023). Several physiological, biochemical, and molecular mechanisms are employed by citrus plants to cope with drought and salinity stress, including the optimization of stomatal regulation and photosynthetic gas exchange, maintenance of cellular osmotic balance, activation of enzymatic and non-enzymatic antioxidant defenses, and dynamic reprogramming of phytohormonal networks involved in stress perception and response (Arbona et al., 2003, 2005; Zandalinas et al., 2016). In this regard, the adoption of stress-tolerant rootstocks constitutes a key agronomic approach to enhance citrus productivity under drought and salinity constraints. Evidence from previous studies indicates marked differences in the adaptive responses of citrus rootstocks to abiotic stress, with Sunki mandarin (*Citrus sunki*) displaying relatively low tolerance to water deficit, while Swingle citrumelo [*C. paradisi* (Macfadyen) x *P. trifoliata* (L.) Raf.] has consistently exhibited greater resilience to both water and salt stress (Modica et al., 2024; Silva et al., 2023). Carrizo citrange (*Citrus sinensis* cv. Washington navel x *Poncirus trifoliata*) is among the most extensively adopted citrus rootstocks; nevertheless, it exhibits high sensitivity to abiotic stress factors, particularly drought and salinity (Modica et al., 2024; Moya et al., 2002; Scialò et al., 2024). In contrast, recent evidence indicates that Bitters (*C. sunki* × *P. trifoliata*) displays enhanced tolerance to PEG-induced water deficit, outperforming Carrizo citrange in terms of physiological and growth responses under drought stress conditions (Scialò et al., 2024). To cope with drought and salinity, citrus plants deploy a wide range of physiological, biochemical, and molecular responses. These include the regulation of stomatal conductance and photosynthetic gas exchange, the maintenance of cellular osmotic balance, and the activation of enzymatic and non-enzymatic antioxidant defense systems (Arbona et al., 2003, 2005). In addition, stress conditions trigger a dynamic reprogramming of phytohormonal networks involved in stress perception, signaling, and adaptive responses (Zandalinas et al., 2016).

The plant microbiome, defined as the ensemble of microorganisms inhabiting different plant compartments and their functions associated to a specific environment, has gained considerable attention for its role in plant health and productivity (Raaijmakers et al. 2009; Vandenkoornhuyse et al. 2015). Microbial communities associated with the root system, including the rhizosphere and the endorhizosphere, can enhance nutrient acquisition, modulate phytohormone levels, and contribute to plant tolerance against both biotic and abiotic stresses (Berendsen et al., 2012; Compant et al., 2019; Mendes et al., 2013). These beneficial effects are often mediated by plant growth-promoting microorganisms (PGPMs), which can regulate ethylene through ACC deaminase activity, produce phytohormones, and improve osmotic balance under stress conditions (Kelbessa et al., 2023; Ma et al., 2020; Mumtaz et al., 2022). Moreover, the mitigation of abiotic stresses through beneficial microbes in agriculture represents a safe and sustainable strategy to enhance crop productivity as climate change-driven reductions in water availability and increasing soil salinity represent major constraints to plant productivity (Phour & Sindhu, 2022), as well as for citrus (Balfagón & Gómez-Cadenas, 2025).

In citrus, the composition and function of the microbiome have been increasingly investigated, with the aim of understanding plant-microbe interactions and developing microbiome-based strategies to address citriculture challenges (Xu et al., 2018; Zhang et al., 2017, 2021). Large-scale studies have revealed the existence of a global citrus core microbiome, characterized by a set of conserved bacterial and fungal taxa consistently associated with citrus roots across diverse geographical locations (Xu et al., 2018). These core taxa are enriched in the rhizosphere compared to bulk soil and are thought to be selectively recruited by the host based on functional traits, supporting the concept of plant-driven microbiome assembly. Despite these advances, most studies on citrus microbiome have focused on biotic stress conditions, including diseases such as Huanglongbing (HLB), Phytophthora root rot and Mal secco disease where microbial community shifts have been associated with disease progression, inoculum sources and/or plant health status (Dimaria et al., 2023; Ginnan et al., 2020; Mosca et al., 2024; Trivedi et al., 2012; Yang & Ancona, 2021). In these pathosystems, a depletion of beneficial microorganisms and a restructuring of the microbiome have been widely reported. In contrast, much less attention has been devoted to the role of the microbiome in citrus responses to abiotic stresses such as drought and salinity, despite growing evidence suggesting that root-associated plant microbial communities can contribute to mitigating these detrimental effects (Caddell et al., 2019; Mumtaz et al., 2022). Recent studies have characterized microbial communities associated with citrus root systems, often including different rootstocks within their experimental frameworks (Castellano-Hinojosa et al., 2023; Dimaria et al., 2023; Leonardi et al., 2026; Lombardo et al., 2024; Long et al., 2025; Padhi et al., 2019; Penyalver et al., 2022; Song et al., 2020). These studies have provided insights into citrus microbiome composition, although in most cases rootstocks were not the primary focus of these studies, and direct comparisons among genotypes remain limited, with only a few comparative analyses available (Long et al., 2025; Padhi et al., 2019; Penyalver et al., 2022). In addition, the interaction between rootstock genotype, abiotic stress, and microbiome assembly remains insufficiently explored and in particular, the extent to which drought and salinity influence genotype-dependent microbial recruitment has not yet been systematically assessed. This represents a critical knowledge gap, considering that rootstocks are the primary determinants of plant responses to drought and salinity in citrus. Recent studies have demonstrated that citrus rootstocks exhibit markedly different physiological and molecular responses to water deficit, with tolerant genotypes such as Bitters showing a more controlled response compared to sensitive ones such as Carrizo (Scialò et al., 2024; Modica et al., 2024). Similarly, differential physiological strategies involving hormonal regulation, antioxidant activity, and osmotic adjustment have been reported among citrus rootstocks exposed to abiotic stress (Modica et al., 2025).

However, whether these genotype-dependent stress responses are associated with distinct patterns of microbiome assembly and microbial recruitment remains largely unexplored. In particular, the interplay between abiotic stress, rootstock genotype, and the structure and function of root-associated microbial communities has not yet been comprehensively investigated in citrus.

In this context, the present study aims to investigate how water deficit and salinity stress influence the assembly of bacterial and fungal communities associated with two citrus rootstocks differing in stress tolerance, Bitters and Carrizo, across rhizosphere and endorhizosphere compartments. By integrating microbiome profiling with plant morphological, physiological and biochemical analyses, this work seeks to disentangle the relative contributions of microhabitat, stress, and genotype in shaping root-associated microbial communities, and to assess whether genotype-dependent stress responses are associated with distinct patterns of microbial recruitment.

## Materials and methods

### Plant material and experimental design

Two citrus rootstocks, Bitters (BIT) and Carrizo Citrange (CRZ), the first recently released by the University of Riverside, California (Caruso et al., 2020), were used in this study. Seeds were extracted from mature fruits and seeded in pots. Seeds were sown into pre-moistened substrate composed by peat, coconut fiber, sand, and perlite (50:25:20:5). After transplant and during growth plants were irrigated and fertilized using a Hoagland solution as modified by Forner-Giner et al. 2011 until the beginning of the experiment. Ten homogeneous one-year-old plants per rootstock were selected for each treatment. Seedlings were subjected to water deficit and salinity stress for 77 days, i.e. plants were fully irrigated twice weekly (control, 100% Et0) or partially (water stress, WS, 50% Et0) as reported by Modica et al. (2024), whereas 10 plants were irrigated adding 60 mM NaCl (salt stress, SS), as reported by Modica et al. (2025) (Supplementary Figure S1). Morphological, physiological and biochemical measurements were performed at the end of the trial. The total leaf chlorophyll content was assessed following the procedure described by Modica et al. (2024). Leaf chlorophyll content (CCM) was monitored using a CCM-200 plus chlorophyll meter on four plants per treatment, measuring two fully expanded leaves per plant. Leaf water potential (Ψ, MPa) was measured using a pressure chamber (PMS 600, PMS Instruments, Corvallis, OR, USA). Measurements were taken on 3 leaves per rootstock and per treatment between 8:00 h and 10:00 h (solar time), as previously reported by Vanella et al., 2023. Abscisic acid (ABA) quantification was carried out on 15 mg freeze-dried leaves, as described by (Balfagón et al., 2022). Proline content was measured spectrophotometrically using a spectrophotometer (NanoDrop 2000, Thermo Scientific, Waltham, MA, USA), according to Modica et al. (2025). Catalase (CAT, EC 1.11.1.6) was determined as described by Aebi (1978).

### Sample preparation, DNA extraction and library preparation for amplicon sequencing

Root samples were processed according to Anzalone et al. 2021, 2022, with minor modifications. Rhizosphere samples were obtained by gently shaking the roots to remove the non-adhering soil particles. Five grams of roots with firmly attached soil were transferred in sterile 50-mL centrifuge tubes containing 20 mL of sterile saline buffer (0.85% NaCl) and then mixed well by vortex for 2 min. Endorhizosphere samples were obtained by surface sterilizing the previously processed roots, which were first submerged in 75% ethanol solution (2 min), followed by 50% sodium hypochlorite solution (2 min), and 75% ethanol solution (1 min), and rinsed five times in sterile distilled water. In order to verify the surface sterility of the samples, surface-sterilized roots were placed on Potato Dextrose Agar (PDA, Oxoid, Milan, Italy) plates at 27 °C ± 2 for 7 days. The lack of bacterial and fungal growth confirmed the sterility of the root surfaces. Endorhizosphere samples (approximately 5 g) were then homogenized with a mortar and pestle in 20 mL of sterile saline buffer (0.85% NaCl). One mL of rhizosphere soil suspension and of endorhizosphere homogenate for each sample was aliquoted into 2-mL reaction tubes and centrifuged at 13,000 rpm and 4 °C for 30 min. Pellets were then stored at −80 °C for further processing. Total genomic DNA was extracted with DNeasy PowerSoil Pro Kit (Qiagen, Hilden, Germany), according to the manufacturer’s instructions. DNA concentration and quality were determined with a NanoDrop 1000 spectrophotometer (Thermo Scientific, Wilmington, DE, USA). Library preparation and amplicon sequencing were performed at IGA Technology Services (I-33100 Udine, Italy). Bacterial community composition was assessed by amplifying the V3–V4 hypervariable region of the 16S rRNA gene using primers 16S-341F and 16S-805R (Klindworth et al., 2013). To minimize host-derived contamination, peptide nucleic acid (PNA) clamps were included during the initial PCR to suppress chloroplast and mitochondrial 16S rRNA amplification. Fungal communities were profiled by targeting the ITS1 region of the rRNA operon with primers ITS1 and ITS2 (White et al., 1990). Both amplicon libraries were sequenced on an Illumina NovaSeq 6000 platform (Illumina, San Diego, CA, USA) using paired-end 250 bp reads.

### Bioinformatic analysis

All analyses of microbial communities were performed in R (v. 4.5.2) using the relevant packages, except for network analyses. Quality filtering, denoising, trimming, merging forward and reverse reads and chimera removal for the ASVs generation have been performed using DADA2 (v. 1.26) (Callahan et al., 2016) in R (4.0.2). The 16S SILVA database (v.138) (Pruesse et al., 2007) for 16S reads and the UNITE database for the ITS reads (version 9 ‘all eukaryotic dynamic’) (Abarenkov et al., 2020) were considered for the taxonomic identification of the bacterial and fungal communities, respectively. Plant-related sequences (e.g. chloroplast and mitochondria) were filtered out from the 16S ASV table. In the ITS ASV table, non-fungal eukaryotic taxa were excluded to focus exclusively on the fungal community. For both 16S and ITS feature tables, ASVs identified as ‘Unassigned’ were removed as well. Alpha- and beta-diversity analyses based on the ASV tables for the bacterial and fungal communities, using the phyloseq package (version 3.17) in R (v. 4.0.2) (McMurdie & Holmes, 2013), were evaluated for each the rhizosphere and endorhizosphere of the control, water-stressed and salinity-stress plants in Bitters and Carrizo rootstocks. Statistical significance was analysed using the Kruskal-Wallis test for alpha-diversity and the PERMANOVA test (999 permutations) for beta-diversity through vegan (v.2.6-4) (Oksanen et al., 2001). Differential abundance was assessed with DESeq2 (v. 1.40.2) (Love et al., 2014) to detect enrichment and depletion of bacterial and fungal genera between treatments and controls within each microhabitat and genotype. Wald test was considered for each bacterial and fungal genus across the comparisons with the parametric fit approach and the resulting *P*-values for each independent statistical test were adjusted according the False Discovery Rate (FDR) method (threshold of individual *P*-values < 0.05). Core microbiome analysis was performed at the genus taxonomic levels for bacterial and fungal communities, considering a prevalence of 99% in the samples for bacterial taxa and 75% for fungal taxa using the microbiome R package (Lahti & Shetty, 2018). Graphical outputs of alpha- and beta-diversity and PCA biplots were obtained through the use of ‘ggplot2’ (v. 4.0.2) (Wickham, 2009). The relative abundances of core bacterial and fungal taxa were visualized as heatmaps using the ‘ComplexHeatmap’ (v.2.26.1) (Z. Gu, 2022) R package. Upset plots representing the differential abundance comparisons were visualized using ‘UpSetR’ (v. 1.4.1) (Conway et al., 2017) R package. Correlation matrices were obtained using ‘corrplot’ (v. 0.95) R package (Wei et al., 2017). Microbial network analyses were performed through ‘MENAP’ (Deng et al., 2012) following the developer’s recommendations and choosing the greedy modularity as separation method. Bacterial and fungal networks were built for each genotype and treatments from both compartments, considering only ASVs with a relative abundance greater than 0.5% and occurring in both compartments in each subset. Gephi (v. 0.9) (Bastian et al., 2009) was used for visualization and the total degree values have been considered to assess the central nodes.

### Statistical analysis

Morphological (root volume), physiological (ABA, chlorophyll content, CCM, and leaf water potential), and biochemical (catalase activity and proline content) parameters were first tested for normality using the Shapiro-Wilk test. Proline content in the BIT_SS group did not meet the assumption of normality and was analyzed separately using the Kruskal-Wallis test, followed by Dunn’s post hoc multiple-comparison test. All other parameters were analyzed using a two-way ANOVA in R with the ‘rstatix’ package (Kassambara, 2019), with treatment and genotype included as fixed factors; statistical significance was set with a *P*-value < 0.05. When significant effects were detected, pairwise comparisons among stress treatments within each genotype were performed using Tukey’s post hoc test, and significance was determined based on *P*-values < 0.05. Correlation analyses were performed using the Pearson’s rho coefficient implemented in the ‘Hmisc’ R package (Harrell Jr et al, 2019) to evaluate associations among phenotypic variables, whose statistical significance was assessed with a *P*-value < 0.05. The correlation of these variables have been tested also with the most significantly enriched bacterial genera in each treatment within the endorhizosphere of both genotypes. Similarly, Principal Component Analysis (PCA) in R was performed to summarize multivariate relationships among phenotypic traits and to visualize their joint distribution with the enriched bacterial genera across treatments using the Facto.

## Results

### Abiotic stress and rootstock drive microhabitat-specific shifts in the citrus root microbial communities

Citrus are composite plants formed by the grafting of a scion onto a rootstock. The rootstock, being the component in direct contact with the soil, is primarily responsible for water and nutrient uptake as well as for the perception of and response to abiotic stresses. To investigate how water (WS) and salt stress (SS) shape the root-associated microbiome we profiled the bacterial and fungal communities of two citrus rootstocks Bitters (BIT) and Carrizo (CRZ) in both the rhizosphere (R) and endorhizosphere (E) compartments, comparing stressed plants to their untreated controls (CK).

After the removal of sequencing primers, adapters, chimera and low quality sequences, a total of 9,729,886 and 2,930,689 16S and ITS reads were retrieved, respectively. The retained bacterial 16S reads, after chloroplast and mitochondria removal, accounted for 68.70% of the total reads, with an average of 172,669 reads per sample. Fungal ITS reads accounted in total for 44.63%, with an average of 27,251 reads per sample.

In order to assess the impact of abiotic stresses on the microbial communities of both rootstocks, alpha-diversity analysis was performed by considering both observed richness and Shannon diversity indexes (Supplementary Figure S2). Overall, these parameters showed statistically significant differences only as an effect of the compartment (Kruskal-Wallis test, *P*-value < 0.05) and were significantly higher in the rhizosphere than in the endorhizosphere.

Principal Coordinate Analysis (PCoA) based on Bray-Curtis distances was performed to visualize the overall structural changes in bacterial and fungal communities across compartments, namely rhizosphere and endorhizosphere, rootstocks, and treatments. In the rhizosphere, bacterial communities were significantly shaped by stress treatments (Fig. 1A). A clear differentiation was observed between treated (WS and SS) and untreated (CK) samples within each rootstock (BIT and CRZ) (PERMANOVA test, R^2^”Treatment”= 0.29, *P-value* < 0.05). Although not reaching statistical significance, a trend toward separation was also noted between BIT_WS and CRZ_WS samples, suggesting a potential rootstock-specific response to water stress in the rhizosphere. In the endorhizosphere, bacterial community structure was influenced by a more complex set of factors. Significant effects based on the interaction between the genotype and treatment (PERMANOVA test R^2^”Genotipe:Treatment” = 0.10, *P-value* < 0.05), indicating that the endophytic bacterial assemblages are shaped not only by the stress itself but also by the specific rootstock genotype and by how each genotype responds to the stress (Fig. 1B). Fungal communities exhibited in the rhizosphere significant differences among stresses and control plants within both rootstocks (PERMANOVA test, R^2^”Treatment” = 0.11, *P*-value < 0.05), closely reflecting the shifts observed in bacterial assemblages (Fig. 1C). In the endorhizosphere, fungal communities were significantly affected by treatment, genotype, and their interaction (PERMANOVA test, R^2^”Genotype:Treatment” = 0.11, *P* value < 0.05) (Fig. 1D), similarly to the bacterial communities in the same compartment.

**Figure 1.**
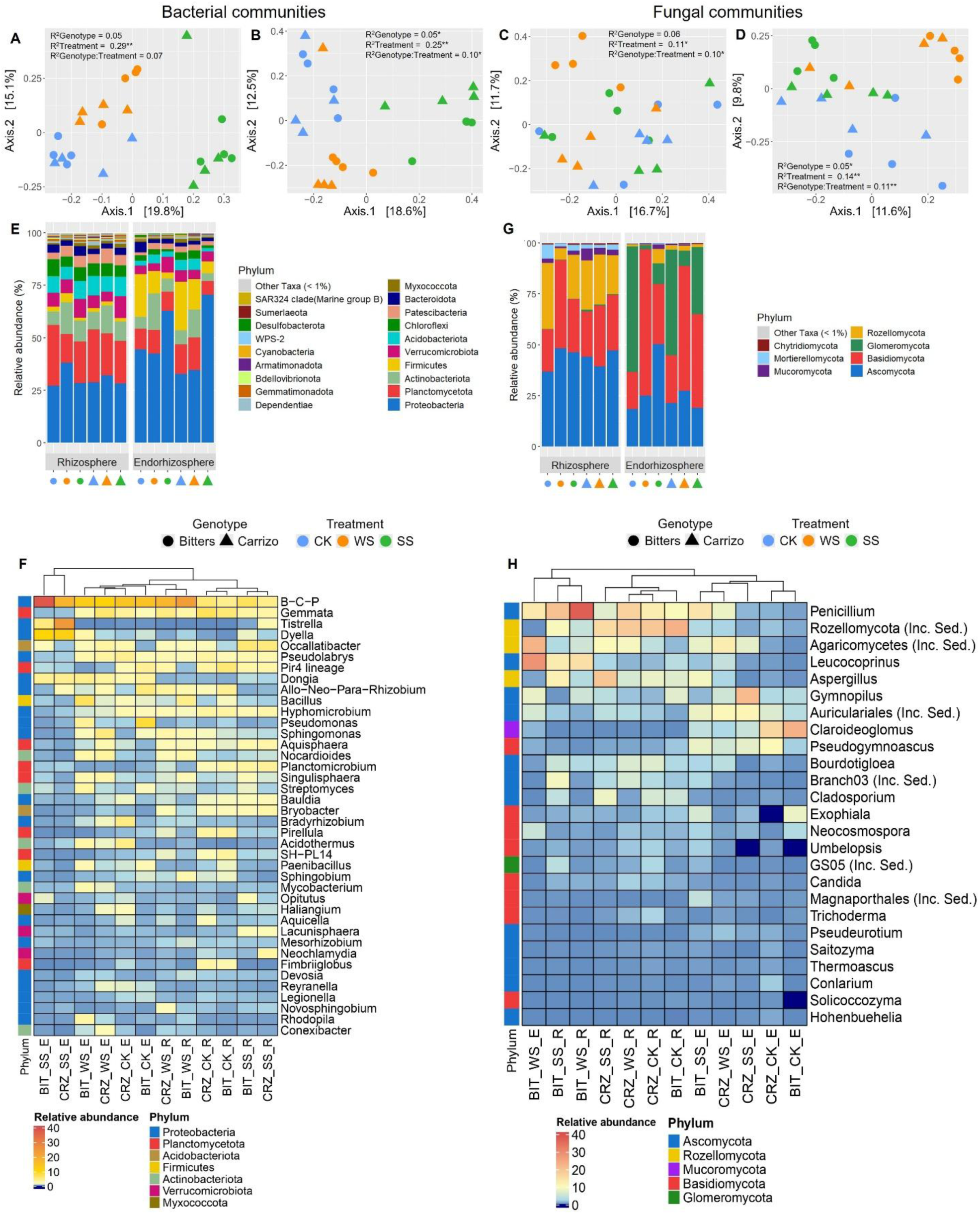
Taxonomic and diversity comparison across compartments, treatments and genotypes. Beta-diversity analysis of the bacterial rhizosphere and endorhizosphere (A,B) bacterial communities (A,B) and (C,D) fungal communities, respectively. The proportion of variation in pairwise distances associated with genotype, treatment and their interaction is reported as R². Statistical significance was determined by PERMANOVA analysis (999 permutations), with one or asterisks indicating *P* value < 0.05 or *P* value ≤ 0.001, respectively. Relative abundance of the (E) bacterial and (G) fungal communities at the phylum taxonomic level; taxa less abundant than 1% are represented as “Other Taxa (< 1%). Relative abundance of the 40 most abundant (F) bacterial core members and all (H) fungal core members detected with a prevalence ≥ 99% and ≥ 75% across the samples, respectively. Acronyms: BIT (Bitters), CRZ (Carrizo); CK (control), WS (water-stress), SS (salinity-stress); R (rhizosphere) and E (endorhizosphere); B-C-P (*Burkholderia-Caballeronia-Paraburkholderia*).

### Treatment- and genotype-specific patterns in the root-associated bacterial communities composition

The most represented phyla of the bacterial communities in the rhizosphere (R) and endorhizosphere (E) of Bitters (BIT) and Carrizo (CRZ) rootstocks under control (CK), water stress (WS), and salt stress (SS) conditions were considered at a relative abundance >1%. Across all samples, six phyla Proteobacteria, Planctomycetota, Actinobacteriota, Verrucomicrobiota, Acidobacteriota, and Patescibacteria accounted together for more than 80% of the relative abundance in each sample (Fig. 1E).

In the rhizosphere, the relative abundance of Proteobacteria and Actinobacteriota increased in the water stress samples of both rootstocks compared to their controls. In BIT, Proteobacteria rose from 27.1% to 38.1% and Actinobacteriota from 6.3% to 15.2%; in CRZ, the increases were from 28.8% to 32.1% and from 8.5% to 13.2%, respectively. Salt stress, in turn, promoted a higher abundance of Actinobacteriota, Bacteroidota, and Patescibacteria in the rhizosphere of both rootstocks. In BITS_SS, their relative abundances increased compared to control plants: Actinobacteriota rose from 8.5% to 9.7%, Bacteroidota from 3.8% to 4.4%, and Patescibacteria from 3.0% to 5.5%. A similar trend was observed in Carrizo, whereby Actinobacteriota increased from 8.5% to 9.3%, Bacteroidota from 2.5% to 3.5%, and Patescibacteria from 4.4% to 5.1%. In the endorhizosphere, the bacterial communities were dominated by Proteobacteria, Firmicutes, Planctomycetota, Actinobacteriota, and Verrucomicrobiota, together representing over 80% of the relative abundance in each sample. Reduction of Verrucomicrobiota was observed under salt stress in both rootstocks (from 20.3% in BIT_CK to 3.0% in BIT_SS; from 23.1% in CRZ_CK to 5.7% in CRZ_SS). Conversely, Proteobacteria increased markedly under salt stress, particularly in CRZ (from 32.7% in CRZ_CK to 70.4% in CRZ_SS), while Actinobacteriota were more abundant under water stress in both genotypes (from 5.8% in BIT_CK to 17.4% in BIT_WS; from 6.8% in CRZ_CK to 13.3% in CRZ_WS).

Bacterial community composition at the genus level was characterized by the presence of dominant taxa consistently distributed across the plant specimens, along with other genera with an overall relative abundance lower than 1%. Core microbiome analysis, defined by a 99% prevalence threshold across the compartments of the plant genotypes, revealed that these bacterial taxa accounted for an average of 62% and 71.54% of the total relative abundance in the rhizosphere and endorhizosphere compartments, respectively. Clear differences in relative abundances were observed between treatments and no-treated specimens within each genotype. This divergence was further confirmed by cluster analysis, which grouped the samples into treatment-specific clusters (Fig. 1F), confirming that stress, rather than genotype, drives the most prominent compositional changes in the rhizosphere. The genus *Burkholderia-Caballeronia-Paraburkholderia* (*B-C-P*) emerged as the dominant taxon under water stress in both rootstocks, increasing markedly in BIT_WS (23.6% vs. 4.3% in BIT_CK) and CRZ_WS (17.5% vs. 5.9% in CRZ_CK). Other genera followed a similar trend, including *Bacillus* (4.5% in BIT_WS vs. 1.6% in BIT_CK; 1.2% in CRZ_WS vs. 0.9% in CRZ_CK) and *Sphingomonas* (2.1% vs. 1.3% in BIT; 2.7% vs. 1.6% in CRZ). Under salt stress, the response was more genotype-specific. In BIT_SS, multiple genera increased relative to BIT_CK, including B-C-P (7.2% vs. 4.4%), *Occallatibacter* (5.8% vs. 1.0%), *Planctomicrobium* (3.2% vs. 2.7%), *Dyella* (1.5% vs. 0.1%), and *Streptomyces* (1.8% vs. 1.0%). In contrast, CRZ_SS exhibited a more selective enrichment, with *Allo-Neo-Para-Rhizobium* showing a dramatic increase from 0.02% to 2.2%, while B-C-P declined relative to CRZ_CK (4.5% vs. 5.9%).

Consistent with the rhizosphere, endorhizosphere bacterial communities differed among treatments within each genotype (Fig. 1F). Unlike the rhizosphere, some taxa within treatment-specific clusters were more abundant only in particular genotypes under water or salinity stress. *B-C-P* remained the most abundant genus across treatments, reaching its highest relative abundance under salt stress in both BIT_SS (38.1%) and CRZ_SS (18.3%). Under water stress, *Sphingomonas* and *Bacillus* increased in both genotypes, with *Sphingomonas* reaching 2.7% in BIT_WS and 3.3% in CRZ_WS, and *Bacillus* reaching 2.3% and 1.5%, respectively. Notably, CRZ_WS showed a distinctive enrichment of Pir4 lineage (6.1% vs. 3.7% in CRZ_CK) and *Bauldia* (5.1% vs. 0.5%). Under salt stress, *Planctomicrobium* and *Occallatibacter* were highly enriched in both rootstocks. *Planctomicrobium* increased from 0.1% to 5.3% in BIT and from 0.8% to 11.6% in CRZ. *Occallatibacter* rose from 0.1% to 5.3% in BIT and from 0.2% to 24.5% in CRZ, becoming the dominant taxon in CRZ_SS. Additional genotype-specific enrichments were observed, including *Singulisphaera* and *Mycobacterium* in BIT_SS (averaging 1.6% vs. <0.5% in controls), and *Hyphomicrobium* (2.0% vs. 1.0%) and *Gemmata* (4.1% vs. 2.4%) in CRZ_SS.

### Fungal community composition across treatments and microhabitats

At the phylum level, fungal communities in the rhizosphere of both rootstocks were mostly represented by Ascomycota, Basidiomycota and Rozellomycota (Fig. 1G). In the rhizosphere of Bitters, both stress treatments increased the relative abundance of the dominant fungal phyla compared to the control. Ascomycota rose from 36.73% in BIT_CK to an increase of 46.34% in BIT_SS while Basidiomycota increased from 20.80% in BIT_CK to 43.46% in BIT_WS. The response in Carrizo (CRZ) was more variable, Ascomycota decreased under water stress (39.33% vs. 44.02% in CRZ_CK), but reached the highest abundance under salt stress (47.17% in CRZ_SS). Basidiomycota, conversely, increased under both stresses, as observed in Bitters (27.33% in CRZ_WS and 29.81% in CRZ_SS) compared to the control (22.37%). In the endorhizosphere, taxonomic shifts were more pronounced. In Bitters, Ascomycota expanded from 18.4% in BIT_CK to 50.4% in BIT_SS, while Basidiomycota dominated under water stress reaching the 71.81% in BIT_WS. Notably, Glomeromycota, which were highly prevalent in the control (61.63%), underwent a drastic reduction under both stresses dropping to 2.12% in BIT_WS and 10.16% in BIT_SS. A similar pattern was observed in Carrizo. Ascomycota decreased in CRZ_SS compared to CRZ_CK, and Basidiomycota abundance fluctuated reaching from 23.39% (CK) to 61.31% in CRZ_WS. Glomeromycota, as in Bitters was most abundant in CRZ_CK (51.84%) decreasing to 7.59% in CRZ_WS.

The fungal core microbiome, defined using a 99% prevalence threshold across all samples, comprised only four taxa. These taxa were the only fungal genera consistently detected in nearly all plant specimens, indicating that the fungal core members were markedly reduced in size compared with the bacterial counterpart. Considering a prevalence threshold of 75%, the fungal core microbiome expanded to 25 members predominantly identified at the genus level and belonging to the phyla Ascomycota, Basidiomycota, Rozellomycota, Mucoromycota, and Glomeromycota. As observed for the bacterial communities, the root compartment (rhizosphere vs. endorhizosphere) was the primary factor differentiating fungal community composition. However, within each compartment, no consistent clustering by treatment was detected across the rootstock, in contrast to what was observed in bacterial communities (Fig. 1G).

In the rhizosphere, several genera showed a similar composition based on their genotypes. In Bitters, *Penicillium* increased under both WS and SS (20.5% and 37.4%, respectively) compared to BIT_CK (9.7%), as *Leucocoprinus* (12.8% and 15.1% vs. 3.5%) (Fig. 1H). *Aspergillus* increased only in BIT_SS (11.6% vs. 3.5% in BIT_CK). In Carrizo, Agaricomycetes incertae sedis increased under both stresses (10.1% in CRZ_WS, 9.2% in CRZ_SS vs. 4.0% in CRZ_CK). *Penicillium* was more abundant in CRZ_WS (17.3%) than in CRZ_CK (12.6%) and CRZ_SS (7.1%), while the highest relative abundance of *Aspergillus* was in CRZ_SS (19.9% vs. 8.1% in CRZ_CK and 6.0% in CRZ_WS). In the endorhizosphere, stress-induced shifts were more pronounced. In Bitters, *Penicillium* was the most abundant genus under both WS and SS (13.6% and 14.0%, respectively), compared to 0.8% in BIT_CK (Fig. 1H). *Leucocoprinus* and Agaricomycetes incertae sedis dominated in BIT_WS (23.0% and 28.4%, respectively), while in BIT_SS, *Aspergillus* and *Pseudogymnoascus* emerged with an average abundances of 8.5%. In Carrizo, the two treatments shared a similar abundance of Auriculariales incertae sedis (10.8% in CRZ_WS, 11.2% in CRZ_SS). However, each treatment also exhibited unique patterns: CRZ_WS showed higher abundances of *Penicillium*, *Leucocoprinus*, and *Aspergillus* (8.3%, 4.4%, and 2.6%, respectively) than CRZ_CK (1.3%, 0.6%, and 0.9%), while CRZ_SS was characterized by a distinct enrichment of *Gymnopilus* (21.4% vs. 2.8% in CRZ_CK).

### Stress-dependent and genotype-specific assembly of the citrus root microbiome

In the rhizosphere, most enriched taxa were treatment-specific and unique to each genotype, despite consistently belonging to the same dominant phyla, Actinobacteriota and Proteobacteria, across all conditions (Fig. 2, see Data Set S1 at https://10.5281/zenodo.20829953). In BIT, the cumulative relative abundance of statistically significant enriched taxa was 38.01% under WS and decreased to 25.11% under SS. A decrease was observed in CRZ whose corresponding values were 4.19% under WS and 10.17% under SS.

**Figure 2.**
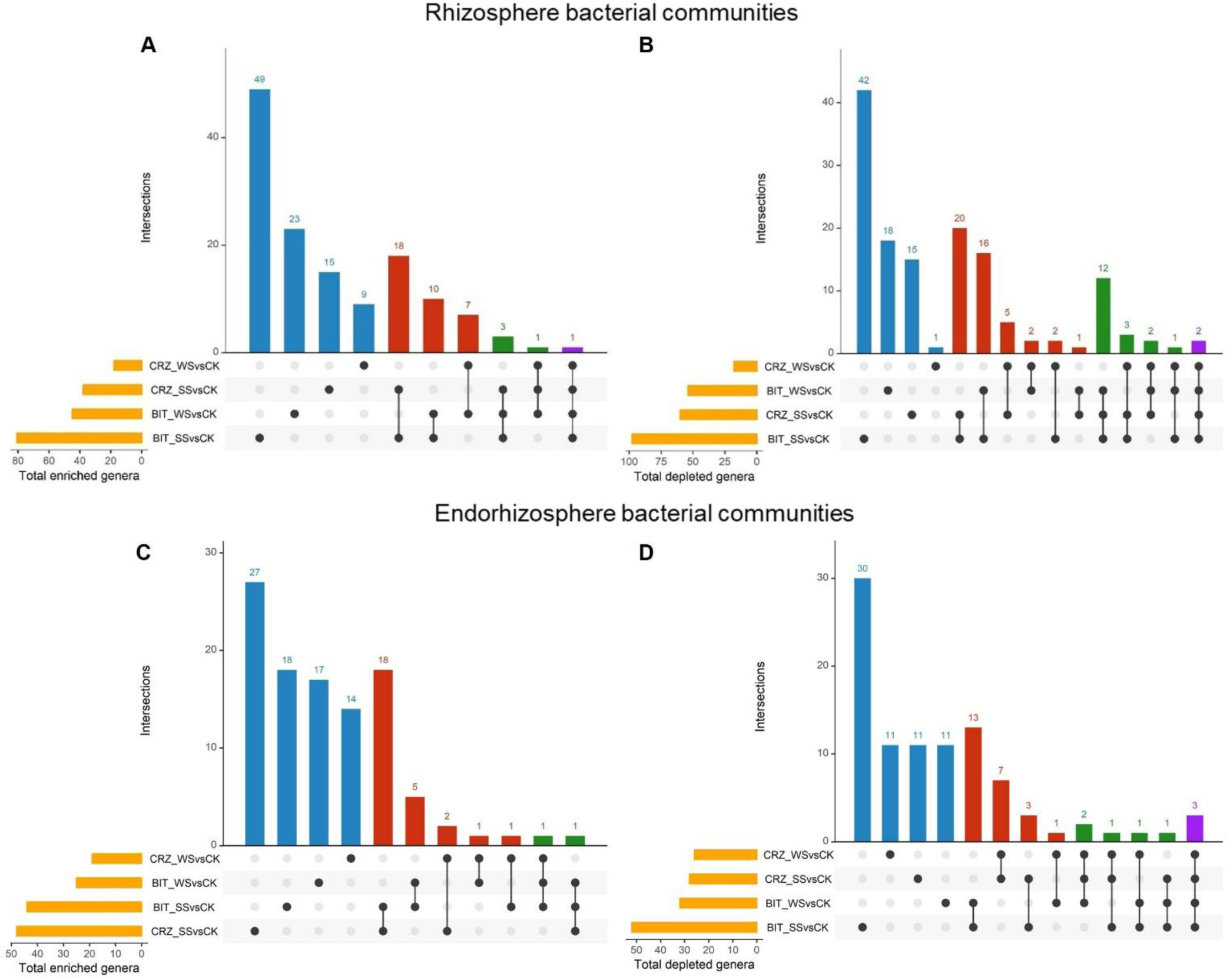
Upset plots showing the number of (A, C) enriched and (B, D) depleted bacterial genera (Wald test, individual *P*-values < 0.05, FDR-corrected) in each treatment relative to their respective untreated controls. Horizontal orange bars denote the total number of differentially enriched or depleted genera per comparison, while vertical bars show the counts of genera that are unique to or shared across the pairwise comparisons. The connected dots below the vertical bars indicate the specific combinations of comparisons that share the same genera, and the bar colours (blue, red, green, and purple) are colour-coded to these combinations.

In BIT_WS vs. BIT_CK 44 genera, mainly represented by Actinobacteriota and, to a lesser extent, Proteobacteria phyla, were enriched, with 23 unique to this treatment, including *Nocardioides, Conexibacter, Knoellia* (Actinobacteriota) and *Nitrobacter, Brevundimonas, Pseudacidovorax, Rhodopila* (Proteobacteria) (Fig. 2A). In BIT_SS vs. BIT_CK, 80 taxa were enriched, with 49 exclusively in this condition. These were predominantly distributed among Proteobacteria (e.g., *Thalassospira Alkanibacter*), Actinobacteriota (e.g*., Salinibacterium, Actinomadura, Nocardia, Sinomonas*) and Bacteroidota (e.g., *Tunicatimonas*, *Flavimarina*, *Persicitalea*, *Rubrivirga*). Moreover, a set of bacterial genera shared by Bitters under both stress conditions was identified, comprising 10 taxa mainly belonging to Proteobacteria such as *Thalassobaculum, Marinobacter, Falsirhodobacter* and core genus *Tistrella*. On the other hand, Carrizo showed no enriched and shared taxa between WS and SS. In CRZ_WS vs. CRZ_CK, nine genera were enriched, belonging to Proteobacteria, Actinobacteriota, and a single Firmicutes taxon, whereas in CRZ_SS vs. CRZ_CK 15 enriched genera were identified, distributedamong Proteobacteria (e.g., *Nitrospirillum*, *Pelagibacterium*, *Hyphomonas*) and Bacteroidota (e.g., *Salegentibacter*, *Vitellibacter*) (Fig. 2A). Cross-genotype comparisons under the same treatment revealed no shared taxa between BIT_WS and CRZ_WS, consistent with beta-diversity results. Conversely, BIT_SS and CRZ_SS shared 19 taxa, predominantly belonging to Proteobacteria (*Alcanivorax*, *Cellvibrio*, *Marinobacter*, *Thalassobaculum*) and Bacteroidota (*Marinoscillum*, *Crocinitomix*). Regarding the depleted taxa (i.e. higher in controls), Proteobacteria was the most representative phylum under WS in both genotypes, whereas a marked depletion of Rhizobiales and Burkholderiales members was observed under SS, suggesting particular sensitivity of these orders to salinity regardless of host genotype.

In the endorhizosphere, bacterial taxa enriched in each treatment relative to their respective controls showed distinct quantitative patterns between the two genotypes. In BIT, the cumulative abundance of enriched taxa was 10.20% under WS and 3.45% under SS. In CRZ, the corresponding values were 12.11% under WS and markedly increased to 54.23% under SS. Moreover, the genera enriched under each treatment predominantly belonged to the same phyla, with a compositional continuity across treatments for each genotype.

In BIT_WS vs. BIT_CK, 24 enriched genera were identified, with 17 unique to this treatment and belonging to Proteobacteria (e.g. *Rhizomicrobium, Nitrobacter*) and Actinobacteriota (*Acidothermus, Nocardia* and *Sinomonas*) (Fig. 2C). In BIT_SS vs. BIT_CK, 43 enriched taxa were found, with 18 unique to this condition such as *Siccibacter, Acidiphilum* and *Rhizomicrobium* (Proteobacteria), *Actinophytocola* (Actinobacteriota) and *Pseudopedobacter* and *Marinoscillum* (Bacteroidota). Five genera were shared between BIT_WS and BIT_SS, with a prevalence of Actinobacteriota taxa, such as *Nocardia, Sinomonas, Frankia* and *Parafrigoribacterium*. In CRZ_WS vs. CRZ_CK, 15 enriched taxa were identified, mostly belonging to Actinobacteriota (*Nocardioides*, *Marmoricola*, *Arthrobacter*) and Proteobacteria (*Sphingomonas*, *Snodgrassella*, *Frischella*) (Fig. 2C). In the comparison between CRZ_SS and CRZ_CK, enriched genera were identified, with a prevalence of Bacteroidota (*Marinoscillum*, *Marinoscillum*, *Salegentibacter*) and Proteobacteria where three enriched genera, namely *Tistrella, Dyella* and *Dongia* were among the most abundant ones and part of the core members, along with *Allorhizobium-Neorhizobium-Pararhizobium-Rhizobium* and *Ferrovibrio* (Fig. 2C). Cross-genotype comparisons revealed only a single bacterial genus under WS conditions (*Acidicapsa*, Actinobacteriota), whereas a shared response under SS was observed between BIT and CRZ, with six genera in common, primarily within Bacteroidota, including the enriched genera *Imperialibacter*, *Marinoscillum*, and *Pseudopedobacter*. Consistent with rhizosphere patterns, the endorhizosphere of BIT_SS showed a significant depletion of taxa belonging to the orders Rhizobiales and Burkholderiales, whereas in the other treatments and genotypes depleted taxa were more heterogeneously distributed across taxonomic groups rather than being concentrated within specific orders (Fig. 2D).

Differential abundance analysis showed that rhizosphere fungal communities contained fewer taxa than bacterial communities (Supplementary Figure S4; see Data Set S1 at https://10.5281/zenodo.20829953). Salinity stress was the main driver of exclusive fungal genera in both rootstocks. In BIT_SS vs. BIT_CK, only *Aureobasidium* (Ascomycota) was enriched, whereas in CRZ_SS vs. CRZ_CK, seven genera, mainly represented by Basidiomycota (e.g., *Volvarella*, *Tulasnella*, *Moesziomyces*) were exclusively enriched (Supplementary Figure S3A). Under water stress, no exclusive taxa were detected in BIT_WS vs. BIT_CK, while only one genus was enriched in CRZ_WS vs. CRZ_CK. Depleted taxa were mainly represented by members of Ascomycota and Basidiomycota, shared between the two treatments of Bitters, whereas Carrizo showed only different groups of no shared taxa (Supplementary Figure S3B). In particular, the depleted taxa in CRZ_WS vs CRZ_CK were represented by six genera in total, five of them represented by Basidiomycota and only one by *Glomus* (Glomeromycota). In CRZ_SS vs CRZ_CK, only two Basidiomycota genera were depleted.

In the endorhizosphere, significant enrichments were observed only in Bitters, and no depleted taxa were detected across treatments in either genotype (Supplementary Figure S3C). Enriched genera belonged exclusively to Basidiomycota and Glomeromycota. Most of them were shared between these two treatments in Bitters (e.g. *Tulasnella*, Basidiomycota; *Dominkia*, Glomeromycota) whereas only *Rhizophagus* (Glomeromycota) was enriched in BIT_WS and two Basdiomycota taxa in BIT_SS (Supplementary Figure S3D).

### Water and salinity stresses differentially shape microbial networks and central bacterial taxa in Bitters and Carrizo rootstocks

To gain further insight into the effect of water and salinity stress on microbial interactions, co-occurrence networks were constructed for bacterial and fungal communities in the rhizosphere and endorhizosphere of both rootstocks, comparing stressed plants to their respective controls (Table 1, Data Set S2 at; see Data Set S2 at https://10.5281/zenodo.20829953). Overall, the networks of the water- and salinity-stressed plants differed across the two genotypes. In Bitters, salinity stress increased network complexity: despite a similar number of nodes across treatments (Table 1), the number of edges was substantially higher in BIT_SS compared to BIT_CK and BIT_WS. Accordingly, the clustering coefficient increased in SS, while modularity decreased. In WS condition, modularity increased, depicting a more compartmentalized network structure.

**Table 1.** Topological properties of control, leaf-inoculated, and root-inoculated co-occurrence bacterial networks in Bitters and Carrizo.

| Kingdom | Genotype | Treatment | No. of nodes | No. of edges | Clustering Coefficient | Modules |
| --- | --- | --- | --- | --- | --- | --- |
| Bacteria | BIT | CK | 230 | 562 | 0.21 | 23 |
|  |  | WS | 177 | 500 | 0.15 | 27 |
|  |  | SS | 210 | 828 | 0.36 | 18 |
|  | CRZ | CK | 279 | 782 | 0.26 | 27 |
|  |  | WS | 226 | 327 | 0.21 | 35 |
|  |  | SS | 217 | 325 | 0.18 | 31 |
| Fungi | BIT | CK | 53 | 591 | 0.45 | 0 |
|  |  | WS | 82 | 1953 | 0.7 | 2 |
|  |  | SS | 88 | 2204 | 0.65 | 1 |
|  | CRZ | CK | 99 | 1934 | 0.47 | 2 |
|  |  | WS | 69 | 271 | 0.19 | 6 |
|  |  | SS | 41 | 265 | 0.23 | 5 |

**Table 2.** Morphological (root volume), physiological (chlorophyll content, chlorophyll content meter [CCM], leaf water potential, abscisic acid [ABA]), and biochemical parameters (proline content and catalase activity) measured in the two citrus rootstocks, Carrizo and Bitters, subjected to water stress (WS, 50% H₂O) and salt stress (SS, 60 mM), compared with the untreated control (CK). Two-way ANOVA was used to assess the effects of genotype, stress treatment, and their interaction. Data are presented as mean ± standard deviation. Within each column, values followed by the same letter are not significantly different according to Tukey’s post hoc test (P < 0.05), with comparisons performed among stress treatments within each genotype. For proline content in the Bitters genotype, statistical significance was assessed using the Kruskal-Wallis test, followed by Dunn’s post hoc multiple-comparison test (*P* value < 0.05) among stress treatments within that genotype.

| Rootstock | Treatments | Root volume (cm<sup>3</sup>) | Chlorophyll (µg/cm<sup>2</sup>) | CCM | Leaf Water potential (MPa) | ABA (ng g<sup>-1</sup> dry weight) | Proline (mmol g<sup>-1</sup>) | Catalase (mmol H<sub>2</sub>O<sub>2</sub>/mg protein min) |
| --- | --- | --- | --- | --- | --- | --- | --- | --- |
| Bitters | CK | 56.00 ± 11.54 a | 15.42 ± 1.28 | 60 ± 14.14 a | -1.50 ± 0.00 a | 8071.32 ± 677.30 a | 387.15 ± 64.65 a | 2.03 ± 1.51 a |
| | WS | 68.4 ± 22.05 ab | 21.18 ± 2.14 | 74.9 ± 7.93 a | -1.43 ± 0.15 a | 7011.38 ± 1112.01 a | 1653.41 ± 365.96 ab | 0.64 ± 0.52 a |
| | SS | 89.39 ± 9.42 b | 12.85 ± 1.44 | 47.1 ± 27.75 a | -1.80 ± 0.20 a | 8072.13 ± 1483.75 a | 6543.77 ± 68.54 b | 1.70 ± 0.49 a |
| Carrizo | CK | 95.67 ± 31.34 a | 15.09 ± 0.44 | 72.3 ± 19.03 a | -1.33 ± 0.12 | 5784.45 ± 704.72 a | 276.32 ± 37.00 | 1.59 ± 0.34 |
| | WS | 58.50 ± 4.50 b | 23.88 ± 1.33 | 95.7 ± 14.21 a | -1.81 ± 0.03 a | 9600.80 ± 1147.10 b | 791.75 ± 42.96 a | 4.68 ± 1.15 a |
| | SS | 71.33 ± 3.05 ab | 17.46 ± 0.75 | 74.63 ± 19.60 a | -1.85 ± 0.48 a | 7260.14 ± 1858.84 ab | 1203.25 ± 396.85 a | 4.75 ± 1.06 a |
| Statistical method | Terms | Root volume (cm<sup>3</sup>) | Chlorophyll (µg/cm<sup>2</sup>) | CCM | Water potential (MPa) | ABA (ng g<sup>-1</sup> dry weight) | Proline (mmol g<sup>-1</sup>) | Catalase (mmol H<sub>2</sub>O<sub>2</sub>/mg protein min) |
| ANOVA/Kruskal-Wallis<sup>1</sup> | Rootstock | ns | 0.005 | 0.01 | ns | 0.7 | <0.001 | <0.001 |
|  | Treatment | ns | <0.0001 | 0.7 | <0.01 | 0.01 | <0.001 | <0.05 |
|  | Interaction | <0.001 | <0.05 | 0.04 | ns | 0.002 | NA<sup>2</sup> | <0.001 |

<sup>1</sup> Statistical significance was evaluated using the Kruskal-Wallis test only for proline.
<sup>2</sup> Interaction effects were not examined for proline, as this parameter was subjected to a non-parametric analysis (Kruskal-Wallis test) for the BIT\_SS treatment.

In the BIT_CK network, the nodes in central positions were mostly represented by ASVs identified in genera such as *Rhodoplanes, Inquilinus, Burkholderia-Caballeronia-Paraburkholderia, Chthonomonas* (Proteobacteria), *Nostoc* (Cyanobacteria) and *Terrimicrobrium* (Firmicutes) (Fig. 3A). The central nodes in BIT_WS were represented by ASVs identified as core genera, such as *B-C-P, Sphingomonas and Defluviicoccus* (Proteobacteria)*, Mycobacterium, Arthrobacter and Kribbella* (Actinobacteriota) (Fig. 3B). Regarding the BIT_SS networks, the central nodes with the highest number of degree were identified within the core genera members *Acidothermus, Paenibacillus, Pseudomonas and Sphingobium* (Fig. 3C). In CRZ, bacterial networks exhibited a different structure compared to BIT. Across the abiotic stresses in comparison to the no-treated CRZ speckimes, CRZ_WS and CRZ_SS networks were characterized by a lower number of edges across all treatments, an increase of modules and node numbers relatively stable (Table 1, Fig. 3D-F). Consequently, the number of central nodes was lower than in Bitters which were mainly represented by three ASVs identified within Proteobacteria and Gemmatimonadota phyla. The network depicting interactions in CRZ_WS, which was the highest modularized, showed as central nodes ASVs identified in genera belonging to Proteobacteria (represented by *Sumerlea*) and Actinobacteriota (*Conexibacter and Iamia*), in common with the network of BIT_WS plants. The other central nodes were represented by ASVs identified as Candidatus *Solibacter* (Acidobacteriota), *Planctopirus*, SH-PL14 (Planctomycetota) and Candidatus *Protochlamydia* (Verrucomicrobiota). Concering the network of Carrizo salinity-stressed plants, also ASVs of Proteobacteria (*Dyella,* Ellin6067*, Roseiarcus* and *Dongia)* and Actinobacteriota (*Gaiella* and *Conexibacter*) genera were shared with the network of Bitters as central nodes. The other central nodes were ASVs identified mostly in Acidobacteriota (*Terracidiphilus,* Candidatus *Solibacter, Bryobacter* and *Granulicella*) and Myxococcota (*Haliangium)* phyla.

**Figure 3.**
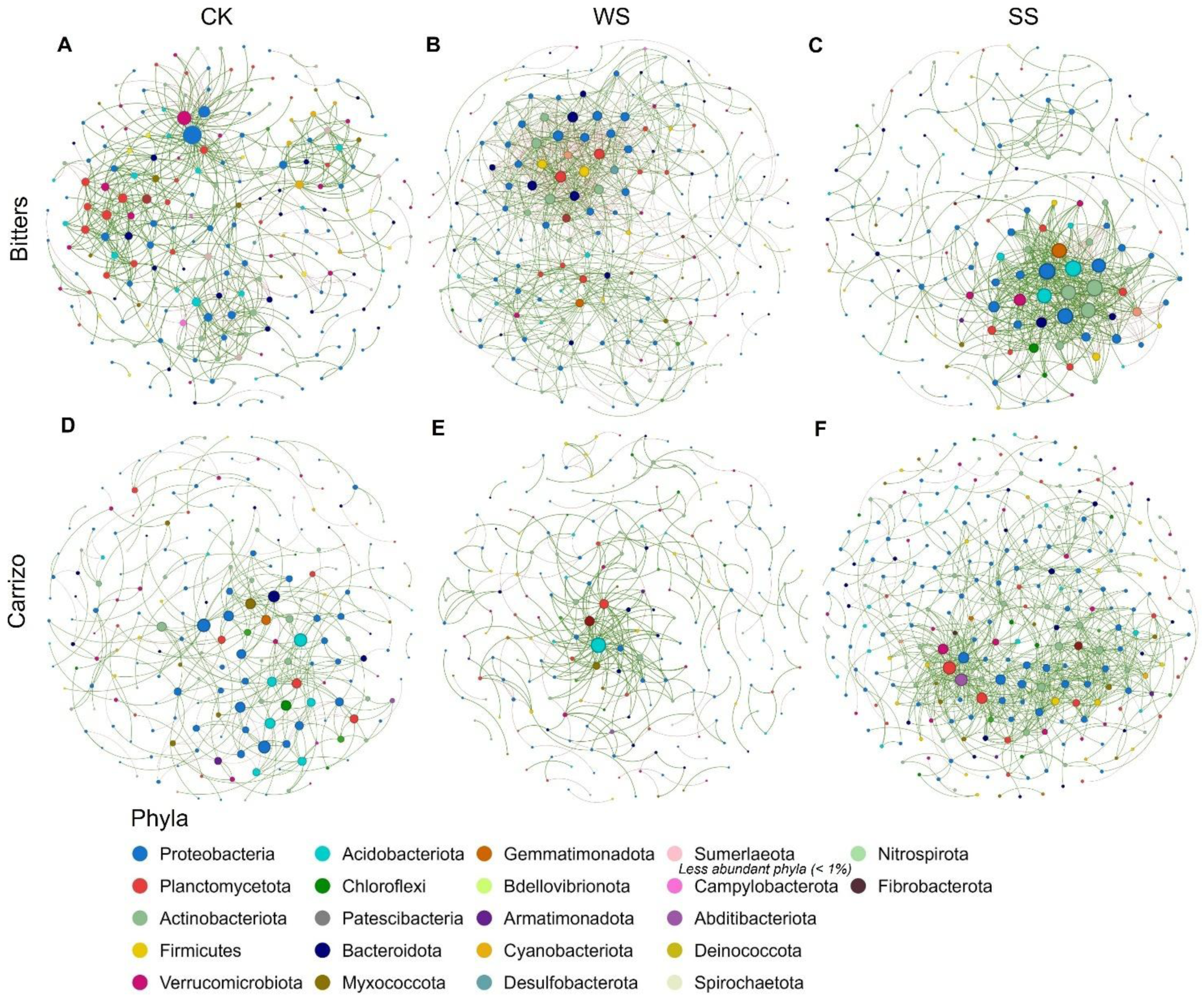
Co-occurrence networks of bacterial communities in the rhizosphere and endorhizosphere of control, water-stressed, and salinity-stressed plants, corresponding to the (A-C) Bitters and (D-F) Carrizo genotypes. Nodes are colored according to the phylum-level classification of the corresponding ASVs, and their size is proportional to node degree. Positive and negative correlations are indicated by green and red edges, respectively.

Fungal networks showed distinct patterns: in BIT_WS and BIT_SS networks a higher number of edges compared to CK was observed, despite a moderate increase in node numbers (Table 1, Supp. Fig. S3). The clustering coefficient was higher under stress than in CK, while the number of modules remained low. In Carrizo, networks depicting the interactions in both stresses showed an opposite trend. Both WS and SS networks showed a marked reduction in nodes and compared to CK (Table 1). However, modularity increased under stress, with more modules in CRZ_WS and CRZ_SS than in CRZ_CK. Due to the absence of a defined network structure, no significant differences in node centrality measures were detected across fungal networks.

### Genotype-specific strategies for water- and salinity-stress tolerance based on plant phenotypes and associated bacterial taxa

Rootstock responses to water and salinity stresses were assessed by integrating morphological, physiological, and biochemical parameters and comparing them with their respective untreated controls (Tab. 1). Root volume, representing the morphological trait, exhibited a highly significant genotype × treatment interaction (two-way ANOVA, *P*-value < 0.001). BIT showed a significant increase in root volume in BIT_WS and BIT_SS relative to BIT_CK, whereas CRZ displayed a significant reduction in CRZ_WS and CRZ_SS compared with CRZ_CK. Physiological parameters related to chlorophyll, CCM, and leaf water potential revealed distinct patterns. CCM values were significantly influenced by the genotype × treatment interaction (p = 0.04). Leaf water potential was significantly affected by treatment (two-way ANOVA, *P* value < 0.01), with CRZ_SS showing the strongest water-deficit effect, reflected in markedly lower values than CRZ_CK. Regarding biochemical traits, ABA levels displayed a significant genotype × treatment interaction and increased substantially in CRZ_WS and CRZ_SS compared with CRZ_CK. Proline content also varied significantly across the genotype × treatment interaction (two-way ANOVA, *P* value < 0.001), with BIT_SS showing the highest accumulation among all groups. Catalase activity was similarly shaped by the genotype × treatment interaction (two-way ANOVA, *P* value < 0.001), with CRZ_WS and CRZ_SS exhibiting markedly higher CAT activity than their respective controls.

These results prompted an investigation of potential correlations among parameters within each genotype, considering both untreated and stressed plants. Pearson’s rho correlation analysis indicated that in BIT_WS and BIT_SS increases in root volume and proline content were positively correlated, whereas BIT_CK showed no such relationship (Fig. 4A-C). In BIT_CK ABA increase correlated positively and significantly with leaf water potential, but not with catalase activity; the latter correlation emerged only in BIT_WS. In CRZ, no significant correlation was detected between root volume and proline content across treatments but, instead, significant correlations were found between catalase activity and leaf water potential across all treatments (Fig. 4D-F). Nevertheless, unlike BIT_WS, no significant association between ABA and leaf water potential was observed in CRZ_WS.

**Figure 4.**
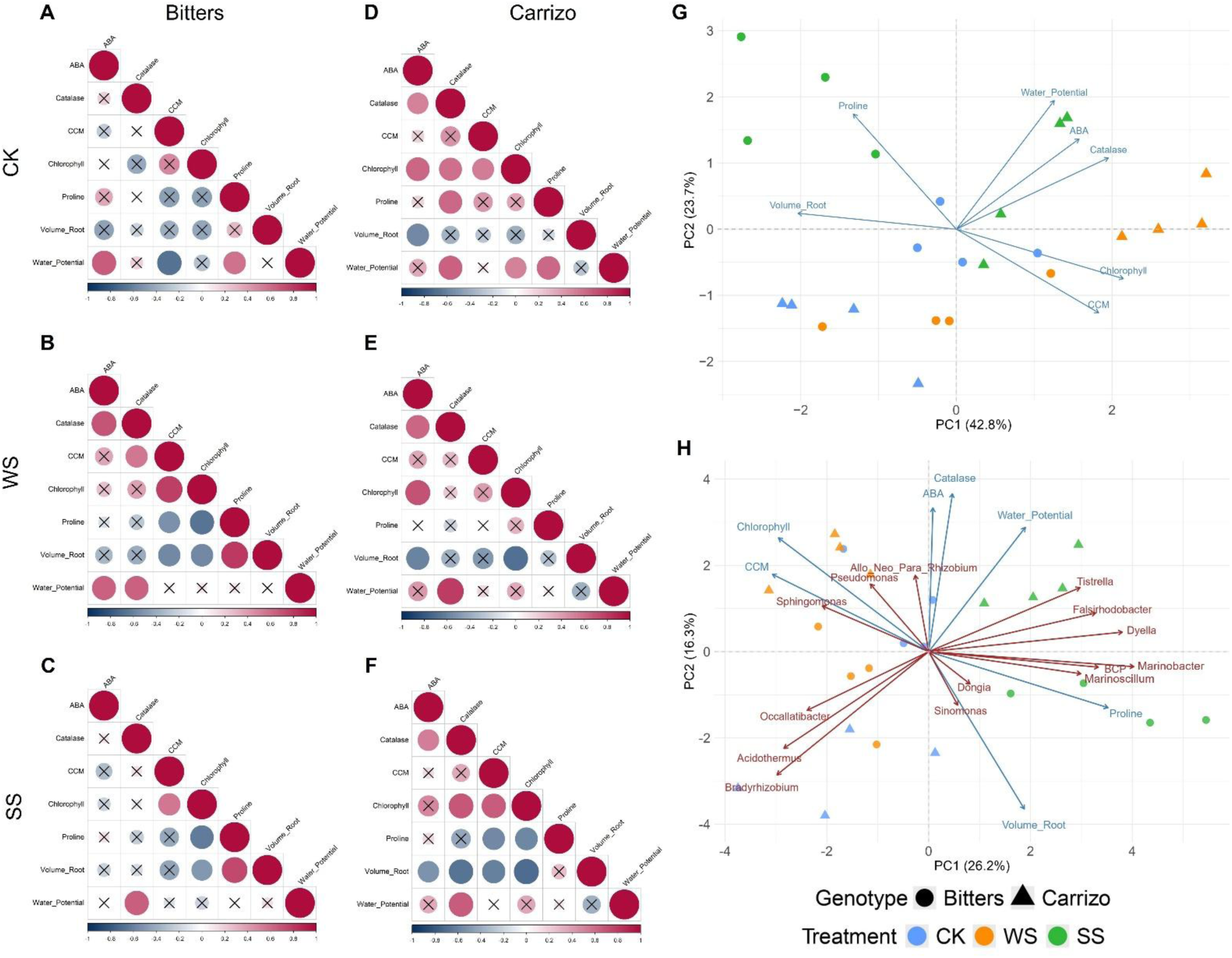
Pearson’s rho correlations among the main physiological, biochemical and morphological parameters in (A-C) Bitters and (D-F) Carrizo under (A,D) control (CK), (B,E) water stress (WS), and (C,F) salt stress (SS) conditions. Circle size is proportional to the magnitude of the correlation coefficient, with red and blue colors indicating positive and negative correlations, respectively. Statistical significance of each comparison was assessed with a *P*-value < 0.05; crosses (×) denote non-significant correlations. (G) Principal Component Analysis (PCA) plot showing the phenotypic variables as loadings, while samples are distinguished by a shape and colour according to genotype and treatment, respectively. (H) PCA plot integrating phenotypic parameters (blue loadings) and specific endorhizosphere bacterial taxa most representative of the core microbiome, network and enrichment analysis in both treatments (red loadings); samples are represented using the same genotype-specific shapes and treatment-specific colours.

Principal component analysis (PCA) revealed a clear discrimination among plant groups based on both rootstock genotype and treatment, collectively accounting for 66.5% of the total variance (Fig. 4G). Specifically, Dim1 discriminated genotypes on the basis of elevated total chlorophyll content, CCM readings, ABA concentration, CAT activity, and leaf water potential, with these latter three variables were predominantly associated with the CRZ rootstock. Conversely, proline accumulation and root volume exhibited stronger positive correlations with the BIT rootstock, suggesting distinct physiological and biochemical adaptive strategies between the two genotypes in response to the imposed treatments.

Relative abundances of the dominant bacterial core members in the endorhizosphere, together with the statistically enriched taxa and the central nodes identified in the co-occurrence networks, were examined in relation to the phenotypic dataset. The two principal components explained 42.5% of the total variance (Fig. 4H). The first principal component (Dim1), separated samples according to stress treatment irrespective of genotype. Within this axis, CRZ_WS plants formed a distinct cluster characterized by elevated chlorophyll and CCM values (Dim1 < 0). These phenotypic traits were positively and significantly correlated with *Sphingomonas* (Person’s rho correlation, individual *P*-value < 0.05). In contrast, BIT_WS samples displayed a more dispersed pattern, suggesting weaker within-group cohesion. The second principal component (Dim2) captured the differentiation between Bitters and Carrizo plants under salinity stress. BIT_SS plants were distinguished by higher abundances of *B-C-P, Marinoscillum, Marinobacter, Dyella, Dongia, and Sinomonas*. These taxa showed a positive and statistically significant correlation with proline content, consistent with the marked proline accumulation observed in BIT_SS plants (Person’s rho correlation, individual *P*-value < 0.05). CRZ_SS plants were instead characterized by low leaf water potential, which correlated with *Tistrella* and *Falsirhodobacter*. Although *Tistrella* was more abundant in CRZ_SS, this core member also showed a positive association with root volume, indicating a broader association with CRZ phenotypic responses.

## Discussion

Water deficit and rising salinity are among the main constraints in Mediterranean environments and citrus production (Lo Piero et al. 2020, Balfagón and Gómez-Cadenas 2025). While the use of rootstocks represents an effective strategy to cope with these adverse conditions, these abiotic stresses still compromise rootstock performance (Modica et al. 2024, 2025) and insights into the associated root microbiome may provide strategies to improve stress tolerance (Caddell et al. 2019). In this study, we investigated how water deficit and salinity affected the composition and structure of bacterial and fungal communities associated with two citrus rootstocks with contrasting water and salinity stress tolerance, integrating microbiome data with plant physiological responses.

Our results show that microbiome composition under abiotic stress is governed by the combined action of compartment, stress, and genotype, with each factor contributing differently. The strongest structuring factor was the root compartment, with a clear differentiation between rhizosphere and endorhizosphere microbial communities. This pattern is consistent with the well-established check-points model of microbiome assembly, according to which the soil acts as a reservoir of microbial diversity, while the plant progressively exerts a stronger selective filter towards internal compartments (Bulgarelli et al., 2013; Reinhold-Hurek et al., 2015). In our study, differentiation of rhizosphere bacterial communities was mainly driven by treatment effects, although other studies on citrus rootstocks have shown that plant genotype can influence the composition of the bacterial communities in response to environmental factors such as water stress (Teng et al., 2026) and amendment applications (Castellano-Hinojosa et al. 2023) also in the rhizosphere microhabitat. Fungal communities displayed more variable responses and were mainly structured by a microhabitat effect, in agreement with previous observations indicating that bacterial communities are generally more responsive to short-term environmental perturbations, whereas fungal assemblages tend to be more stable and habitat-dependent in terms of composition (Barnard et al., 2013) and interactions (de Vries et al., 2018; Gao et al., 2022).

Beyond microhabitat and stress effects, a central finding of this study is the role of rootstock genotype in modulating microbiome assembly under abiotic stress. The two rootstocks exhibited contrasting responses, with Bitters showing broader enrichment genera and increased microbial network complexity, especially under salinity, whereas Carrizo displayed a more constrained response. These differences are consistent with previous physiological and molecular studies on the same genotypes, where Bitters showed more controlled response to water stress, while Carrizo activates stronger oxidative and transcriptional stress responses (Scialò et al., 2024). Such contrasting strategies indicate that the two rootstocks differ not only in stress tolerance, but also in how stress signals are processed and translated into metabolic responses. These differences are likely to influence root-associated microbial communities through changes in root exudation patterns; in addition, plant growth-promoting microorganisms are known to enhance stress tolerance through the production of phytohormones, osmoprotectants, and other bioactive compounds (Ma et al., 2020; Mumtaz et al., 2022). In this context, the more controlled physiological response of Bitters may promote a more stable and selective exudation pattern, favouring a coordinated microbial recruitment, whereas the stronger perturbation observed in Carrizo may result in a less structured microbiome. Consistent with this interpretation, our results provide clear evidence of selective microbial recruitment under stress.

In this study, microbiome shifts were primarily driven by changes in the relative abundance of existing taxa rather than by the complete replacement of community members, supporting that stress induces a reorganization of resident microbiome. Similar patterns have been reported in the citrus rhizosphere microbiome (Castellano-Hinojosa et al, 2023), where enrichment of specific functional groups represents the dominant response to environmental stress. Within this taxonomic framework, Actinobacteriota showed a consistent increase under both water deficit and salinity stress conditions, supporting their role as key components of stress-adapted microbial communities. Members of this phylum are well known for possessing plant-beneficial traits, including phytohormone production, nutrient mobilization, and enhanced tolerance to environmental stressors (Ebrahimi-Zarandi et al., 2023). Their functional relevance in citrus systems is further supported by experimental evidence showing their capacity to promote plant growth and improve nutrient acquisition (Giassi et al., 2016). In the endorhizosphere, fungal communities were dominated by Glomeromycota, particularly under control conditions and salinity stress. This group includes arbuscular mycorrhizal fungi (AMF), which are widely recognized as key symbionts contributing to plant tolerance to drought and salinity through improved nutrient uptake and modulation of stress responses (Gupta et al., 2019; Song et al., 2020); their persistence across treatments suggests a stable and functionally important association with the host plant.

Concerning the most abundant bacterial genera, the top thirty accounted for approximately 75% of the overall relative abundance and were consistently detected across all treatments, compartments, and genotypes. This prompted the analysis of a core microbiome defined at a 99% prevalence threshold, under which none of these dominant taxa were excluded. This observation indicates that the differences observed among treatments and genotypes are not primarily driven by the presence or absence of specific taxa, but rather by shifts in their relative abundances. This pattern is consistent with previous studies on citrus rootstocks, where variations in microbial diversity were reported without major changes in overall community composition (Castellano-Hinojosa et al, 2023). Furthermore, it aligns with evidence demonstrating that root metabolic profiles differ among citrus rootstocks, influencing both the quantity and composition of root exudates (Albrecht et al., 2018, 2019; Dong et al., 2016). It is plausible that genotype-dependent differences in root metabolic profiles, and their modulation under stress, contribute to shaping microbial communities through changes in root exudation, thereby altering the relative abundance of functionally relevant taxa rather than driving taxonomic turnover. The bacterial core microbiome identified in this study was largely composed of taxa affiliated with the dominant phyla of the citrus root microbiome, including Proteobacteria, Planctomycetota, Actinobacteriota, Verrucomicrobiota, and Firmicutes, with only a minor contribution from Myxococcota. A similar taxonomic structure has been previously reported in citrus rootstocks derived from *Poncirus trifoliata* and *Citrus sunki* lineages, which are closely related to Bitters and Carrizo (Penyalver et al., 2022), indicating a conserved phylogenetic framework of the citrus core microbiome. Several core genera previously identified in citrus rhizosphere communities were consistently detected, including *Burkholderia-Caballeronia-Paraburkholderia* (*B-C-P*), *Cellvibrio*, *Devosia, Dyella*, *Mesorhizobium*, *Novosphingobium*, *Pseudomonas*, *Sphingobium*, and *Variovorax* (Xu et al., 2018). Among these, *B-C-P*, *Mesorhizobium*, *Sphingobium*, and *Sphingomonas* showed higher abundance under water stress conditions, particularly in Bitters, suggesting a stress-driven enrichment of taxa with known plant growth-promoting functions. Species belonging to these genera are widely recognized for their role in stress mitigation, due to their ability to produce different volatile organic compounds, phytohormones, enhance nutrient availability regulating plant stress responses (Asaf et al., 2020; Boss et al., 2022; Camacho et al., 2025; Y. Luo et al., 2020). In parallel, genera such as *Bacillus* and *Streptomyces* increased in abundance under water stress, although showing genotype-specific patterns, with *Bacillus* being more represented in Bitters and *Streptomyces* in Carrizo. These taxa are commonly associated with drought tolerance, through mechanisms such as phytohormone production, osmotic adjustment, and modulation of ethylene levels (Lahlali et al., 2022; Yandigeri et al., 2012). Under salinity stress, the rhizosphere communities of the two genotypes showed a more pronounced divergence in core composition. Several genera, including *Occallatibacter*, *Lacunisphaera*, *Aquisphaera*, *Bryobacter*, and *Bauldia* were enriched under saline conditions. Although their functional role in citrus is still poorly understood, some of these taxa have been reported in stress-related or saline environments, suggesting a potential involvement in osmotic adaptation processes (Hagen et al., 2024; J. Luo et al., 2023; Talwar et al., 2020). In contrast to the rhizosphere, the endorhizosphere displayed a stronger selective enrichment of specific core members under stress conditions. Moreover, the applied stresses moderately enriched genera with a relative abundance above 1%, while rare taxa (i.e. those below 1%) were more commonly detected in treated samples, where their presence strongly indicated stress conditions. Under water deficit, genera such as *Streptomyces*, *Novosphingobium*, and *Bacillus* were consistently enriched in both genotypes, confirming their role as common endophytic members of the citrus root microbiome (Zhang et al., 2021). These bacteria are well known for their ability to promote plant growth under drought conditions through IAA production, ACC deaminase activity, and modulation of stress-related metabolic pathways (Aamir et al., 2026; Goudjal et al., 2013). Salinity stress further highlighted the role of specific core taxa in the endorhizosphere, with the *Burkholderia-Caballeronia-Paraburkholderia* complex emerging as one of the most abundant bacterial members, particularly in Bitters. This group has been previously identified as a key component of the citrus root microbiome (Lombardo et al., 2024) and is known to preferentially colonize endosphere and rhizoplane niches (Zhang et al., 2017). In our study, *B-C-P* showed a compartment-dependent pattern, with higher abundance in the rhizosphere under water stress and in the endorhizosphere under salinity stress, suggesting a context-specific ecological role. Several strains belonging to this group have been reported to confer plant tolerance to salinity through mechanisms such as ACC deaminase activity, IAA production, phosphate solubilization, and induction of proline accumulation (Hwang et al., 2025; Maqsood et al., 2021; Zhu et al., 2021).

These core microbiome patterns were consistently reflected in the differential enrichment analysis. Across both genotypes, the greatest number of taxa under stress belonged to the dominant phyla Proteobacteria, Actinobacteriota, and Bacteroidota, with enrichment patterns driven mainly by changes in relative abundance rather than the introduction of new taxa. Notably, rare taxa were frequently detected under stress conditions, suggesting that the rare biosphere may act as a functional reservoir activated under environmental perturbations (Jousset et al., 2017; Pascoal et al., 2021). However, their specific contribution to plant fitness remains to be clarified (Trivedi et al., 2020). At the rhizosphere level, *B-C-P* and *Nocardioides* were among the most enriched genera under water stress, particularly in Bitters. Their role in auxin production, siderophore synthesis, and nutrient mobilization has been previously demonstrated in drought conditions (Ahmad et al., 2022; Kabir et al., 2025; Tuo et al., 2015). In Carrizo, genera such as *Occallatibacter* displayed variable responses, being associated both with non-stress and stress conditions in different studies (Faist et al., 2023; Hagen et al., 2024), highlighting the complexity of their ecological role. Under salinity stress, several taxa were specifically enriched in Carrizo (e.g. *Rheinheimera, Tistrella, Cellvibrio*), although their role in salt stress remains largely unexplored. At the same time, both genotypes shared the recruitment of taxa adapted to saline environments, indicating the existence of a conserved stress-driven selection mechanism.

The response of bacterial co-occurrence networks to water and salt stress was contingent upon rootstock genotype, as evidenced by comparisons with their respective control conditions. In the case of BIT_SS, both the number of nodes and edges increased under treatment, a pattern that was observed in bacterial microbiomes associated with salinity-tolerant grapevine cultivars (B. Wang et al., 2023). In contrast, water deficit induced divergent responses between genotypes: while BIT_WS maintained node and edge counts comparable to those of BIT_CK, CRZ_WS exhibited a higher modularity than CRZ_CK, indicative of a substantial structural disruption. This finding is consistent with the results reported by Gao et al. 2022 and may be attributable to the concurrent marked decreases in root volume observed in this genotype under stress.

The morphological, physiological, and biochemical responses of the different rootstocks to water deficit and salinity stress highlighted a strong dependence on the interaction between genotype and treatment, reflecting distinct adaptive strategies. Root volume was significantly influenced by this interaction, in agreement with previous studies showing that plant responses to salinity are modulated by multiple factors, including stress duration, rootstock genotype, and developmental stage (Alvarez-Gerding et al., 2015; Aparicio-Durán et al., 2021; Othman et al., 2023). Accordingly, Carrizo citrange samples exhibited enhanced sensitivity to salinity, with a reduction of their root volume, confirming the effect of this stress in determining a high sensitivity in this rootstock (Modica et al., 2024; Vives-Peris et al., 2023). At the physiological level, leaf water potential was markedly reduced under both drought and salinity stress, particularly in CRZ_WS and salinity-treated plants (BIT_SS and CRZ_SS), indicating impaired plant water status (Colmenero-Flores et al., 2020). This was accompanied by significant changes in abscisic acid (ABA) levels, with CRZ_WS showing the highest ABA accumulation and suggesting a stronger activation of stress-responsive signaling pathways (Zandalinas et al., 2016). In parallel, proline accumulation was the highest in BIT_SS, pointing to an enhanced osmoregulatory response under salinity. As a multifunctional osmoprotectant, proline contributes to osmotic adjustment, ROS scavenging, and the stabilization of cellular structures (Ghosh et al., 2022). Consistently, CAT activity was also modulated by genotype and treatment, with CRZ plants showing increased activity under stress, likely reflecting enhanced oxidative pressure and the need for antioxidant defense (Alia et al., 2001; Sofo et al., 2015).

Correlation analyses reinforced these patterns, indicating that the Bitters genotype exhibits a more coordinated tolerance to both stress conditions. A significant three-way association among ABA, catalase, and leaf water potential reflects a rapid and integrated response to water stress. Under salinity stress, the strong positive correlation between proline content and root volume further supports Bitters as a tolerant genotype. In contrast, the Carrizo genotype showed a consistent correlation between catalase and ABA across treatments reflecting its higher susceptibility to the imposed water and salt stress conditions and suggesting a tighter coordination between antioxidant defense and ABA-mediated stress signaling (Modica et al., 2024; 2025).

Within this physiological framework, the observed microbiome shifts under salinity provide additional insights into potential biotic contributions to stress adaptation. Taxa such as *Sinomonas*, *Marinobacter*, and *Marinoscillum* increased in abundance under saline conditions and were significantly associated with proline accumulation, suggesting a possible functional link between microbial recruitment and host osmotic regulation. These genera are known to possess plant-beneficial traits and to contribute to osmolyte-related processes (J. Gu et al., 2007; Teng et al., 2026; Y. Wang et al., 2025), raising the possibility that their enrichment may be connected to the enhanced proline accumulation observed, particularly in BIT_SS. Similar associations detected for the core genera (e.g. *Dyella*, *Tistrella*, *Dongia*) further support the hypothesis that microbial mechanisms related to osmolyte biosynthesis and uptake, including proline and ectoine production, may contribute to shaping plant biochemical responses under stress.

Interestingly, both genotypes shared part of this microbial recruitment pattern despite their contrasting physiological responses, suggesting the existence of a conserved microbiome response to salinity superimposed on genotype-specific modulation. This partial convergence may indicate that certain microbial taxa are consistently selected under osmotic stress conditions, regardless of host sensitivity, potentially contributing to baseline stress mitigation processes. In this context, the identification of taxa consistently enriched across genotypes is particularly relevant for the development of synthetic microbial communities (SynComs), as these microorganisms may represent key functional components underpinning osmotic adjustment and stress tolerance. More broadly, core microbiome members are thought to reflect long-term co-evolutionary processes and to contribute to plant fitness through conserved functional traits (Lundberg et al., 2012; Toju et al., 2018), thereby representing promising targets for microbiome-based strategies aimed at improving crop resilience under abiotic stress (Liu et al., 2025).

While this study provides initial insights into the root environment of the microbiome of Bitters and Carrizo under water and saline stress, some aspects may deserve further investigation. The experiment was conducted under controlled pot conditions, which, although useful for reducing environmental variability, may only partially reflect field complexity. In addition, the temporal dimension of microbiome responses across different stress intensities was not extensively explored and could be addressed in future work. The integration of complementary omics approaches may also help to better contextualize the observed microbial patterns from a functional perspective, as well as extending the analysis to a broader range of rootstocks for a more comprehensive understanding of genotype-dependent responses.

## Data Availability

The original data presented in the study are openly available in Zenodo at the following doi: https://10.5281/zenodo.20829953 (accessed on June 25, 2026).

The codes to reproduce data analyses are available at https://github.com/AlexandrosMosca/Microbiome_Citrus_Roostocks

## References

Aamir, M., Shah, K., Moharana, D. P., Validov, S. Z., & Ansari, W. A. (2026). Microbial consortia and drought tolerance-A paradigm shift towards agro-ecological sustainability. World Journal of Microbiology and Biotechnology 2026 42:2, 42(2), 72-. 10.1007/S11274-025-04753-5

Abarenkov, K., Zirk, A., Piirmann, T., Pöhönen, R., Ivanov, F., Nilsson, R. H., & Kõljalg, U. (2020). UNITE QIIME release for Fungi. 10.1093/nar/gkad1039

Aebi, H. F. (1978). Catalase. Methods in Enzymatic Analysis. https://cir.nii.ac.jp/crid/1573950400003728000

Ahmad, H. M., Fiaz, S., Hafeez, S., Zahra, S., Shah, A. N., Gul, B., Aziz, O., Mahmood-Ur-Rahman, Fakhar, A., Rafique, M., Chen, Y., Yang, S. H., & Wang, X. (2022). Plant Growth-Promoting Rhizobacteria Eliminate the Effect of Drought Stress in Plants: A Review. Frontiers in Plant Science, 13, 875774. 10.3389/FPLS.2022.875774/FULL

Albrecht, U., Tripathi, I., & Bowman, K. D. (2019). Rootstock influences the metabolic response to Candidatus Liberibacter asiaticus in grafted sweet orange trees. Trees 2019 34:2, 34(2), 405–431. 10.1007/S00468-019-01925-3

Albrecht, U., Tripathi, I., Kim, H., & Bowman, K. D. (2018). Rootstock effects on metabolite composition in leaves and roots of young navel orange (Citrus sinensis L. Osbeck) and pummelo (C. grandis L. Osbeck) trees. Trees 2018 33:1, 33(1), 243–265. 10.1007/S00468-018-1773-1

Alia, Mohanty, P., & Matysik, J. (2001). Effect of proline on the production of singlet oxygen. Amino Acids, 21(2), 195–200. 10.1007/S007260170026/METRICS

Alvarez-Gerding, X., Espinoza, C., Inostroza-Blancheteau, C., & Arce-Johnson, P. (2015). Molecular and physiological changes in response to salt stress in Citrus macrophylla W plants overexpressing Arabidopsis CBF3/DREB1A. Plant Physiology and Biochemistry, 92, 71–80. 10.1016/J.PLAPHY.2015.04.005

Anzalone, A., Di Guardo, M., Bella, P., Ghadamgahi, F., Dimaria, G., Zago, R., Cirvilleri, G., & Catara, V. (2021). Bioprospecting of Beneficial Bacteria Traits Associated With Tomato Root in Greenhouse Environment Reveals That Sampling Sites Impact More Than the Root Compartment. Frontiers in Plant Science, 12. 10.3389/fpls.2021.637582

Anzalone, A., Mosca, A., Dimaria, G., Nicotra, D., Tessitori, M., Privitera, G. F., Pulvirenti, A., Leonardi, C., & Catara, V. (2022). Soil and Soilless Tomato Cultivation Promote Different Microbial Communities That Provide New Models for Future Crop Interventions. International Journal of Molecular Sciences, 23(15). 10.3390/IJMS23158820

Aparicio-Durán, L., Hervalejo, A., Calero-Velázquez, R., Arjona-López, J. M., Arenas-Arenas, F. J., Teresa, M., Arenas, L., Fernández, J. A., & Garcia-Caparros, P. (2021). Salinity Effect on Plant Physiological and Nutritional Parameters of New Huanglongbing Disease-Tolerant Citrus Rootstocks. Agronomy 2021, Vol. 11, Page 653, 11(4), 653. 10.3390/AGRONOMY11040653

Arbona, V., Flors, V., Jacas, J., García-Agustín, P., & Gómez-Cadenas, A. (2003). Enzymatic and Non-enzymatic Antioxidant Responses of Carrizo citrange, a Salt-Sensitive Citrus Rootstock, to Different Levels of Salinity. Plant and Cell Physiology, 44(4), 388–394. 10.1093/PCP/PCG059

Arbona, V., Iglesias, D. J., Jacas, J., Primo-Millo, E., Talon, M., & Gómez-Cadenas, A. (2005). Hydrogel substrate amendment alleviates drought effects on young citrus plants. Plant and Soil 2005 270:1, 270(1), 73–82. 10.1007/S11104-004-1160-0

Asaf, S., Numan, M., Khan, A. L., & Al-Harrasi, A. (2020). Sphingomonas: from diversity and genomics to functional role in environmental remediation and plant growth. Critical Reviews in Biotechnology, 40(2), 138–152. 10.1080/07388551.2019.1709793

Balfagón, D., & Gómez-Cadenas, A. (2025). Combined abiotic stresses in mediterranean citrus orchards: Impacts, adaptation strategies, and climate-resilient solutions. Scientia Horticulturae, 353, 114462. 10.1016/J.SCIENTA.2025.114462

Balfagón, D., Rambla, J. L., Granell, A., Arbona, V., & Gómez-Cadenas, A. (2022). Grafting improves tolerance to combined drought and heat stresses by modifying metabolism in citrus scion. Environmental and Experimental Botany, 195, 104793. 10.1016/J.ENVEXPBOT.2022.104793

Barnard, R. L., Osborne, C. A., & Firestone, M. K. (2013). Responses of soil bacterial and fungal communities to extreme desiccation and rewetting. The ISME Journal, 7(11), 2229–2241. 10.1038/ISMEJ.2013.104

Bastian, M., Heymann, S., & Jacomy, M. (2009). Gephi: an open source software for exploring and manipulating networks. Proceedings of the International AAAI Conference on Web and Social Media, 3(1), 361–362.

Berendsen, R. L., Pieterse, C. M. J., & Bakker, P. A. H. M. (2012). The rhizosphere microbiome and plant health. Trends in Plant Science, 17(8), 478–486. 10.1016/J.TPLANTS.2012.04.001

Boss, B. L., Wanees, A. E., Zaslow, S. J., Normile, T. G., & Izquierdo, J. A. (2022). Comparative genomics of the plant-growth promoting bacterium Sphingobium sp. strain AEW4 isolated from the rhizosphere of the beachgrass Ammophila breviligulata. BMC Genomics 2022 23:1, 23(1), 508-. 10.1186/S12864-022-08738-8

Bulgarelli, D., Schlaeppi, K., Spaepen, S., Van Themaat, E. V. L., & Schulze-Lefert, P. (2013). Structure and functions of the bacterial microbiota of plants. Annual Review of Plant Biology, 64, 807–838. 10.1146/ANNUREV-ARPLANT-050312-120106

Caddell, D. F., Deng, S., & Coleman-Derr, D. (2019). Role of the plant root microbiome in abiotic stress tolerance. Seed Endophytes: Biology and Biotechnology, 273–311. 10.1007/978-3-030-10504-4_14

Callahan, B. J., McMurdie, P. J., Rosen, M. J., Han, A. W., Johnson, A. J. A., & Holmes, S. P. (2016). DADA2: High-resolution sample inference from Illumina amplicon data. Nature Methods 2016 13:7, 13(7), 581–583. 10.1038/nmeth.3869

Camacho, M., Vaccaro, F., Brun, P., Ollero, F. J., Pérez-Montaño, F., Negussu, M., Martinelli, F., Mengoni, A., Rodriguez-Navarro, D. N., & Fagorzi, C. (2025). Selection and Characterisation of Elite Mesorhizobium spp. Strains That Mitigate the Impact of Drought Stress on Chickpea. Agriculture (Switzerland), 15(15), 1694. 10.3390/AGRICULTURE15151694/S1

Caruso, M., Continella, A., Modica, G., Pannitteri, C., Russo, R., Salonia, F., Arlotta, C., Gentile, A., & Russo, G. (2020). Rootstocks influence yield precocity, productivity, and pre-harvest fruit drop of mandared pigmented mandarin. Agronomy, 10(9), 1305.

Castellano-Hinojosa, A., Albrecht, U., & Strauss, S. L. (2023). Interactions between rootstocks and compost influence the active rhizosphere bacterial communities in citrus. Microbiome 2023 11:1, 11(1), 79-. 10.1186/S40168-023-01524-Y

Colmenero-Flores, J. M., Arbona, V., Morillon, R., & Gómez-Cadenas, A. (2020). Salinity and water deficit. The Genus Citrus, 291–309. 10.1016/B978-0-12-812163-4.00014-0

Compant, S., Samad, A., Faist, H., & Sessitsch, A. (2019). A review on the plant microbiome: Ecology, functions, and emerging trends in microbial application. Journal of Advanced Research, 19, 29–37. 10.1016/J.JARE.2019.03.004

Conway, J. R., Lex, A., & Gehlenborg, N. (2017). UpSetR: an R package for the visualization of intersecting sets and their properties. Bioinformatics, 33(18), 2938–2940. 10.1093/bioinformatics/btx364

de Vries, F. T., Griffiths, R. I., Bailey, M., Craig, H., Girlanda, M., Gweon, H. S., Hallin, S., Kaisermann, A., Keith, A. M., Kretzschmar, M., Lemanceau, P., Lumini, E., Mason, K. E., Oliver, A., Ostle, N., Prosser, J. I., Thion, C., Thomson, B., & Bardgett, R. D. (2018). Soil bacterial networks are less stable under drought than fungal networks. Nature Communications 2018 9:1, 9(1), 3033-. 10.1038/s41467-018-05516-7

Deng, Y., Jiang, Y. H., Yang, Y., He, Z., Luo, F., & Zhou, J. (2012). Molecular ecological network analyses. BMC Bioinformatics, 13(1). 10.1186/1471-2105-13-113

Dimaria, G., Mosca, A., Anzalone, A., Paradiso, G., Nicotra, D., Pritivera, G. F., Pulvirenti, A., & Catara, V. (2023). Sour Orange Microbiome Is Affected by Infections of Plenodomus tracheiphilus Causal Agent of Citrus Mal Secco Disease. Agronomy, 13(3), 654. https://www.mdpi.com/2073-4395/13/3/654

Dong, X., Liu, G., Wu, X., Lu, X., Yan, L., Muhammad, R., Shah, A., Wu, L., & Jiang, C. (2016). Different metabolite profile and metabolic pathway with leaves and roots in response to boron deficiency at the initial stage of citrus rootstock growth. Plant Physiology and Biochemistry, 108, 121–131. 10.1016/J.PLAPHY.2016.07.007

Ebrahimi-Zarandi, M., Etesami, H., & Glick, B. R. (2023). Fostering plant resilience to drought with Actinobacteria: Unveiling perennial allies in drought stress tolerance. Plant Stress, 10, 100242. 10.1016/J.STRESS.2023.100242

Faist, H., Trognitz, F., Antonielli, L., Symanczik, S., White, P. J., & Sessitsch, A. (2023). Potato root-associated microbiomes adapt to combined water and nutrient limitation and have a plant genotype-specific role for plant stress mitigation. Environmental Microbiome 2023 18:1, 18(1), 18-. 10.1186/S40793-023-00469-X

Forner-Giner, M. Á., Rodríguez-Gamir, J., Primo-Millo, E., & Iglesias, D. J. (2011). Hydraulic and Chemical Responses of Citrus Seedlings to Drought and Osmotic Stress. Journal of Plant Growth Regulation 2011 30:3, 30(3), 353–366. 10.1007/S00344-011-9197-9

Gao, C., Xu, L., Montoya, L., Madera, M., Hollingsworth, J., Chen, L., Purdom, E., Singan, V., Vogel, J., Hutmacher, R. B., Dahlberg, J. A., Coleman-Derr, D., Lemaux, P. G., & Taylor, J. W. (2022). Co-occurrence networks reveal more complexity than community composition in resistance and resilience of microbial communities. Nature Communications 2022 13:1, 13(1), 3867-. 10.1038/s41467-022-31343-y

García-Sánchez, F., Jifon, J. L., Carvajal, M., & Syvertsen, J. P. (2002). Gas exchange, chlorophyll and nutrient contents in relation to Na+ and Cl− accumulation in ‘Sunburst’ mandarin grafted on different rootstocks. Plant Science, 162(5), 705–712. 10.1016/S0168-9452(02)00010-9

Ghosh, U. K., Islam, M. N., Siddiqui, M. N., Cao, X., & Khan, M. A. R. (2022). Proline, a multifaceted signalling molecule in plant responses to abiotic stress: understanding the physiological mechanisms. Plant Biology, 24(2), 227–239. 10.1111/PLB.13363

Giassi, V., Kiritani, C., & Kupper, K. C. (2016). Bacteria as growth-promoting agents for citrus rootstocks. Microbiological Research, 190, 46–54. 10.1016/J.MICRES.2015.12.006

Ginnan, N. A., Dang, T., Bodaghi, S., Ruegger, P. M., McCollum, G., England, G., Vidalakis, G., Borneman, J., Rolshausen, P. E., & Caroline Roper, M. (2020). Disease-induced microbial shifts in citrus indicate microbiome-derived responses to huanglongbing across the disease severity spectrum. Phytobiomes Journal, 4(4), 375–387. 10.1094/pbiomes-04-20-0027-R

Goudjal, Y., Toumatia, O., Sabaou, N., Barakate, M., Mathieu, F., & Zitouni, A. (2013). Endophytic actinomycetes from spontaneous plants of Algerian Sahara: indole-3-acetic acid production and tomato plants growth promoting activity. World Journal of Microbiology and Biotechnology 2013 29:10, 29(10), 1821–1829. 10.1007/S11274-013-1344-Y

Gu, J., Cai, H., Yu, S. L., Qu, R., Yin, B., Guo, Y. F., Zhao, J. Y., & Wu, X. L. (2007). Marinobacter gudaonensis sp. nov., isolated from an oil-polluted saline soil in a Chinese oilfield. International Journal of Systematic and Evolutionary Microbiology, 57(2), 250–254. 10.1099/ijs.0.64522-0

Gu, Z. (2022). Complex heatmap visualization. IMeta, 1(3), e43. 10.1002/imt2.43

Gupta, M. M., Chourasiya, D., & Sharma, M. P. (2019). Diversity of arbuscular mycorrhizal fungi in relation to sustainable plant production systems. Microbial Diversity in Ecosystem Sustainability and Biotechnological Applications: Volume 2. Soil & Agroecosystems, 167–186. 10.1007/978-981-13-8487-5_7

Hagen, M., Dass, R., Westhues, C., Blom, J., Schultheiss, S. J., & Patz, S. (2024). Interpretable machine learning decodes soil microbiome’s response to drought stress. Environmental Microbiome 2024 19:1, 19(1), 35-. 10.1186/S40793-024-00578-1

Harrell Jr, F. E., & Harrell Jr, M. F. E. (2019). Package ‘hmisc.’ CRAN2018, 2019, 235–236.

Hwang, H. H., Huang, Y. T., Chien, P. R., Huang, F. C., Wu, C. L., Chen, L. Y., Hung, S. H. W., Pan, I. C., & Huang, C. C. (2025). A plant endophytic bacterium Burkholderia seminalis strain 869T2 increases plant growth under salt stress by affecting several phytohormone response pathways. Botanical Studies 2025 66:1, 66(1), 7-. 10.1186/S40529-025-00453-3

Jousset, A., Bienhold, C., Chatzinotas, A., Gallien, L., Gobet, A., Kurm, V., Küsel, K., Rillig, M. C., Rivett, D. W., Salles, J. F., Van Der Heijden, M. G. A., Youssef, N. H., Zhang, X., Wei, Z., & Hol, G. W. H. (2017). Where less may be more: how the rare biosphere pulls ecosystems strings. The ISME Journal, 11(4), 853–862. 10.1038/ISMEJ.2016.174

Kabir, A. H., Brailey-Crane, P., Abdelrahman, M., Legeay, J., Ahmed, B., Tran, L. S. P., & Bennetzen, J. L. (2025). Ferritin-Mediated Iron Homeostasis and Bacterial Shifts Are Associated With Drought Adaptation in Sorghum. Physiologia Plantarum, 177(4), e70388. 10.1111/PPL.70388;PAGE:STRING:ARTICLE/CHAPTER

Kassambara, A. (2019). rstatix: Pipe-Friendly Framework for Basic Statistical Tests. CRAN: Contributed Packages. 10.32614/CRAN.PACKAGE.RSTATIX

Kelbessa, B. G., Dubey, M., Catara, V., Ghadamgahi, F., Ortiz, R., & Vetukuri, R. R. (2023). Potential of plant growth-promoting rhizobacteria to improve crop productivity and adaptation to a changing climate. CAB Reviews: Perspectives in Agriculture, Veterinary Science, Nutrition and Natural Resources, 2023(2023). 10.1079/cabireviews.2023.0001

Klindworth, A., Pruesse, E., Schweer, T., Peplies, J., Quast, C., Horn, M., & Glöckner, F. O. (2013). Evaluation of general 16S ribosomal RNA gene PCR primers for classical and next-generation sequencing-based diversity studies. Nucleic Acids Research, 41(1), e1–e1.

Lahlali, R., Ezrari, S., Radouane, N., Belabess, Z., Jiang, Y., Mokrini, F., Tahiri, A., Peng, G., Lahlali, R., Tahiri, A, Ezrari, S., Radouane, N, Belabess, Z., Jiang, Y., Mokrini, F., & Peng, G. (2022). Bacillus spp.- Mediated Drought Stress Tolerance in Plants: Current and Future Prospects. 487–518. 10.1007/978-3-030-85465-2_21

Lahti, L., & Shetty, S. (2018). Introduction to the microbiome R package. Preprint at https://microbiome.github.io/Tutorials.

Leonardi, G. R., Mosca, A., Nicotra, D., Massimino, M. E., Dimaria, G., Privitera, G. F., Vitale, A., Polizzi, G., Aiello, D., & Catara, V. (2026). Non-Target Effects of Trichoderma- and Bacillus-Based Products on the Citrus Microbiome. Horticulturae, 12(5), 529. 10.3390/HORTICULTURAE12050529

Liu, S., Wu, J., Cheng, Z., Wang, H., Jin, Z., Zhang, X., Zhang, D., & Xie, J. (2025). Microbe-mediated stress resistance in plants: the roles played by core and stress-specific microbiota. Microbiome 2025 13:1, 13(1), 111-. 10.1186/S40168-025-02103-Z

Lo Piero, A. R. (2020). Abiotic Stress Resistance. 225–243. 10.1007/978-3-030-15308-3_13

Lombardo, M. F., Zhang, Y., Xu, J., Trivedi, P., Zhang, P., Riera, N., Li, L., Wang, Y., Liu, X., Fan, G., Tang, J., Coletta-Filho, H. D., Cubero, J., Deng, X., Ancona, V., Lu, Z., Zhong, B., Roper, M. C., Capote, N., … Wang, N. (2024). Global citrus root microbiota unravels assembly cues and core members. Frontiers in Microbiology, 15, 1405751. 10.3389/FMICB.2024.1405751

Long, C., Fu, X., Wu, Q., Wang, S., Zhou, X., Mao, J., Guo, L., Shi, W., Yang, H., Yang, T., Du, Y., Yue, J., Wu, D., & Liu, H. (2025). Poncirus trifoliata vs. Citrus junos rootstocks: reshaping lemon rhizosphere microecology through microbial and metabolic reprogramming. Frontiers in Microbiology, 16, 1650631. 10.3389/FMICB.2025.1650631

Love, M., Anders, S., & Huber, W. (2014). Differential analysis of count data–the DESeq2 package. Genome Biol, 15(550), 10–1186.

Lundberg, D. S., Lebeis, S. L., Paredes, S. H., Yourstone, S., Gehring, J., Malfatti, S., Tremblay, J., Engelbrektson, A., Kunin, V., Rio, T. G. Del, Edgar, R. C., Eickhorst, T., Ley, R. E., Hugenholtz, P., Tringe, S. G., & Dangl, J. L. (2012). Defining the core Arabidopsis thaliana root microbiome. Nature 2012 488:7409, 488(7409), 86–90. 10.1038/nature11237

Luo, J., Liu, T., Diao, F., Hao, B., Zhang, Z. C., Hou, Y., & Guo, W. (2023). Shift in rhizospheric and endophytic microbial communities of dominant plants around Sunit Alkaline Lake. Science of The Total Environment, 867, 161503. 10.1016/J.SCITOTENV.2023.161503

Luo, Y., Zhou, M., Zhao, Q., Wang, F., Gao, J., Sheng, H., & An, L. (2020). Complete genome sequence of Sphingomonas sp. Cra20, a drought resistant and plant growth promoting rhizobacteria. Genomics, 112(5), 3648–3657. 10.1016/J.YGENO.2020.04.013

Ma, Y., Dias, M. C., & Freitas, H. (2020). Drought and Salinity Stress Responses and Microbe-Induced Tolerance in Plants. Frontiers in Plant Science, 11(13), 591911. 10.3389/FPLS.2020.591911

Maas, E. V. (1993). Salinity and citriculture. Tree Physiology, 12(2), 195–216. 10.1093/TREEPHYS/12.2.195

Maqsood, A., Shahid, M., Hussain, S., Mahmood, F., Azeem, F., Tahir, M., Ahmed, T., Noman, M., Manzoor, I., & Basit, F. (2021). Root colonizing Burkholderia sp. AQ12 enhanced rice growth and upregulated tillering-responsive genes in rice. Applied Soil Ecology, 157, 103769. 10.1016/J.APSOIL.2020.103769

McMurdie, P. J., & Holmes, S. (2013). Phyloseq: An R Package for Reproducible Interactive Analysis and Graphics of Microbiome Census Data. PLoS ONE, 8(4). 10.1371/journal.pone.0061217

Mendes, R., Garbeva, P., & Raaijmakers, J. M. (2013). The rhizosphere microbiome: Significance of plant beneficial, plant pathogenic, and human pathogenic microorganisms. FEMS Microbiology Reviews, 37(5), 634–663. 10.1111/1574-6976.12028

Modica, G., Arcidiacono, F., La Malfa, S., Gentile, A., & Continella, A. (2025). Physiological Response to Foliar Application of Antitranspirant on Avocado Trees (Persea americana) in a Mediterranean Environment. Horticulturae, 11(8), 928. 10.3390/HORTICULTURAE11080928/S1

Modica, G., Di Guardo, M., Puglisi, I., Baglieri, A., Fortuna, S., Arcidiacono, F., Costantino, D., La Malfa, S., Gentile, A., Arbona, V., & Continella, A. (2024). Novel and widely spread citrus rootstocks behavior in response to salt stress. Environmental and Experimental Botany, 225, 105835. 10.1016/J.ENVEXPBOT.2024.105835

Mosca, A., Dimaria, G., Nicotra, D., Modica, F., Massimino, M. E., Catara, A. F., Scuderi, G., Russo, M., & Catara, V. (2024). Soil Microbial Communities in Lemon Orchards Affected by Citrus Mal Secco Disease. Genes, 15(7), 824. 10.3390/genes15070824

Moya, J. L., Tadeo, F. R., Gómez-Cadenas, A., Primo-Millo, E., & Talón, M. (2002). Transmissible salt tolerance traits identified through reciprocal grafts between sensitive Carrizo and tolerant Cleopatra citrus genotypes. Journal of Plant Physiology, 159(9), 991–998. 10.1078/0176-1617-00728

Mumtaz, M. Z., Ahmad, M., Mehmood, K., Sheikh, A. S., Malik, A., Hussain, A., Nadeem, S. M., & Zahir, Z. A. (2022). Role of Plant Growth-Promoting Rhizobacteria in Combating Abiotic and Biotic Stresses in Plants. Microorganisms for Sustainability, 33, 43–104. 10.1007/978-981-16-4843-4_2/

Oksanen, J., Simpson, G. L., Blanchet, F. G., Kindt, R., Legendre, P., Minchin, P. R., O’hara, R. B., Solymos, P., Stevens, M. H. H., & Szoecs, E. (2001). Vegan: community ecology package. (No Title).

Othman, Y. A., Hani, M. B., Ayad, J. Y., & St Hilaire, R. (2023). Salinity level influenced morpho-physiology and nutrient uptake of young citrus rootstocks. Heliyon, 9(2), e13336. 10.1016/j.heliyon.2023.e13336

Padhi, E. M. T., Maharaj, N., Lin, S. Y., Mishchuk, D. O., Chin, E., Godfrey, K., Foster, E., Polek, M., Leveau, J. H. J., & Slupsky, C. M. (2019). Metabolome and microbiome signatures in the roots of citrus affected by huanglongbing. Phytopathology, 109(12), 2022–2032. 10.1094/PHYTO-03-19-0103-R

Pascoal, F., Costa, R., & Magalhães, C. (2021). The microbial rare biosphere: current concepts, methods and ecological principles. FEMS Microbiology Ecology, 97(1), 227. 10.1093/FEMSEC/FIAA227

Penyalver, R., Roesch, L. F. W., Piquer-Salcedo, J. E., Forner-Giner, M. A., & Alguacil, M. del M. (2022). From the bacterial citrus microbiome to the selection of potentially host-beneficial microbes. New Biotechnology, 70, 116–128. 10.1016/J.NBT.2022.06.002

Phour, M., & Sindhu, S. S. (2022). Mitigating abiotic stress: microbiome engineering for improving agricultural production and environmental sustainability. Planta 2022 256:5, 256(5), 85-. 10.1007/S00425-022-03997-X

Pruesse, E., Quast, C., Knittel, K., Fuchs, B. M., Ludwig, W., Peplies, J., & Glöckner, F. O. (2007). SILVA: A comprehensive online resource for quality checked and aligned ribosomal RNA sequence data compatible with ARB. Nucleic Acids Research, 35(21), 7188–7196. 10.1093/nar/gkm864

Raaijmakers, J. M., Paulitz, T. C., Steinberg, C., Alabouvette, C., & Moënne-Loccoz, Y. (2009). The rhizosphere: A playground and battlefield for soilborne pathogens and beneficial microorganisms. Plant and Soil, 321(1–2), 341–361. 10.1007/S11104-008-9568-6

Reinhold-Hurek, B., Bünger, W., Burbano, C. S., Sabale, M., & Hurek, T. (2015). Roots Shaping Their Microbiome: Global Hotspots for Microbial Activity. Annual Review of Phytopathology, 53(Volume 53, 2015), 403–424. 10.1146/ANNUREV-PHYTO-082712-102342/1

Scialò, E., Sicilia, A., Continella, A., Gentile, A., & Lo Piero, A. R. (2024). Transcriptome Profiling and Weighted Gene Correlation Network Analysis Reveal Hub Genes and Pathways Involved in the Response to Polyethylene-Glycol-Induced Drought Stress of Two Citrus Rootstocks. Biology, 13(8), 595. 10.3390/biology13080595

Silva, S. F., Miranda, M. T., Cunha, C. P., Domingues-Jr, A. P., Aricetti, J. A., Caldana, C., Machado, E. C., & Ribeiro, R. V. (2023). Metabolic profiling of drought tolerance: Revealing how citrus rootstocks modulate plant metabolism under varying water availability. Environmental and Experimental Botany, 206, 105169. 10.1016/J.ENVEXPBOT.2022.105169

Sofo, A., Scopa, A., Nuzzaci, M., & Vitti, A. (2015). Ascorbate Peroxidase and Catalase Activities and Their Genetic Regulation in Plants Subjected to Drought and Salinity Stresses. International Journal of Molecular Sciences 2015, Vol. 16, Pages 13561-13578, 16(6), 13561–13578. 10.3390/IJMS160613561

Song, F., Bai, F., Wang, J., Wu, L., Jiang, Y., & Pan, Z. (2020). Influence of Citrus Scion/Rootstock Genotypes on Arbuscular Mycorrhizal Community Composition under Controlled Environment Condition. Plants 2020, Vol. 9, Page 901, 9(7), 901. 10.3390/PLANTS9070901

Talwar, C., Nagar, S., Kumar, R., Scaria, J., Lal, R., & Negi, R. K. (2020). Defining the Environmental Adaptations of Genus Devosia: Insights into its Expansive Short Peptide Transport System and Positively Selected Genes. Scientific Reports 2020 10:1, 10(1), 1151-. 10.1038/s41598-020-58163-8

Teng, Y., Yin, C., Xu, F., Chen, J., Wu, Q., Ye, M., Liu, Y., & Zhu, K. (2026). Citrus Genotype Modulates Rhizosphere Microbiome Structure and Function Under Drought Stress. Plants, 15(1), 77. 10.3390/plants15010077

Toju, H., Peay, K. G., Yamamichi, M., Narisawa, K., Hiruma, K., Naito, K., Fukuda, S., Ushio, M., Nakaoka, S., Onoda, Y., Yoshida, K., Schlaeppi, K., Bai, Y., Sugiura, R., Ichihashi, Y., Minamisawa, K., & Kiers, E. T. (2018). Core microbiomes for sustainable agroecosystems. Nature Plants 2018 4:5, 4(5), 247–257. 10.1038/s41477-018-0139-4

Trivedi, P., He, Z., Van Nostrand, J. D., Albrigo, G., Zhou, J., & Wang, N. (2012). Huanglongbing alters the structure and functional diversity of microbial communities associated with citrus rhizosphere. ISME Journal, 6(2), 363–383. 10.1038/ismej.2011.100

Trivedi, P., Leach, J. E., Tringe, S. G., Sa, T., & Singh, B. K. (2020). Plant–microbiome interactions: from community assembly to plant health. Nature Reviews Microbiology 2020 18:11, 18(11), 607–621. 10.1038/s41579-020-0412-1

Tuo, L., Dong, Y. P., Habden, X., Liu, J. M., Guo, L., Liu, X. F., Chen, L., Jiang, Z. K., Liu, S. W., Zhang, Y. Bin, Zhang, Y. Q., & Sun, C. H. (2015). Nocardioides deserti sp. nov., an actinobacterium isolated from desert soil. International Journal of Systematic and Evolutionary Microbiology, 65(Pt 5), 1604–1610. 10.1099/IJS.0.000147

Vandenkoornhuyse, P., Quaiser, A., Duhamel, M., Le Van, A., & Dufresne, A. (2015). The importance of the microbiome of the plant holobiont. New Phytologist, 206(4), 1196–1206. 10.1111/NPH.13312

Vanella, D., Consoli, S., Continella, A., Chinnici, G., Milani, M., Cirelli, G. L., D’Amico, M., Maesano, G., Gentile, A., La Spada, P., Scollo, F., Modica, G., Siracusa, L., Longo-Minnolo, G., & Barbagallo, S. (2023). Environmental and Agro-Economic Sustainability of Olive Orchards Irrigated with Reclaimed Water under Deficit Irrigation. Sustainability (Switzerland), 15(20), 15101. 10.3390/su152015101

Vives-Peris, V., López-Climent, M. F., Moliner-Sabater, M., Gómez-Cadenas, A., & Pérez-Clemente, R. M. (2023). Morphological, physiological, and molecular scion traits are determinant for salt-stress tolerance of grafted citrus plants. Frontiers in Plant Science, 14, 1145625. 10.3389/fpls.2023.1145625

Wang, B., Wang, X., Wang, Z., Zhu, K., & Wu, W. (2023). Comparative metagenomic analysis reveals rhizosphere microbial community composition and functions help protect grapevines against salt stress. Frontiers in Microbiology, 14, 1102547. 10.3389/fmicb.2023.1102547

Wang, Y., Bin Peng, Zhao, S., Zhou, J., Hazaisi Hanipa, Tian, C., Wang, Y., Peng, · B, Zhao, · S, Tian, C., Zhao, S., Hanipa, · H, & Zhou, J. (2025). Salinity stress reveals keystone metabolites linking rhizosphere metabolomes and microbiomes in Halophyte Suaeda salsa. Plant and Soil 2025 514:1, 514(1), 1219–1239. 10.1007/S11104-025-07457-9

Wei, T., Simko, V., Levy, M., Xie, Y., Jin, Y., & Zemla, J. (2017). Package ‘corrplot.’ Statistician, 56(316), e24.

White, T. J., Bruns, T., Lee, S., & Taylor, J. (1990). Amplification and direct sequencing of fungal ribosomal RNA genes for phylogenetics. PCR Protocols: A Guide to Methods and Applications, 18(1), 315–322.

Wickham, H. (2009). ggplot2: Elegant Graphics for Data Analysis. Springer-Verlag New York. In Media (Vol. 35, Number July).

Xu, J., Zhang, Y., Zhang, P., Trivedi, P., Riera, N., Wang, Y., Liu, X., Fan, G., Tang, J., Coletta-Filho, H. D., Cubero, J., Deng, X., Ancona, V., Lu, Z., Zhong, B., Roper, M. C., Capote, N., Catara, V., Pietersen, G., … Wang, N. (2018). The structure and function of the global citrus rhizosphere microbiome. Nature Communications, 9(1). 10.1038/s41467-018-07343-2

Yandigeri, M. S., Meena, K. K., Singh, D., Malviya, N., Singh, D. P., Solanki, M. K., Yadav, A. K., & Arora, D. K. (2012). Drought-tolerant endophytic actinobacteria promote growth of wheat (Triticum aestivum) under water stress conditions. Plant Growth Regulation 2012 68:3, 68(3), 411–420. 10.1007/S10725-012-9730-2

Yang, C., & Ancona, V. (2021). Metagenomic Analysis Reveals Reduced Beneficial Microorganism Associations in Roots of Foot-Rot-Affected Citrus Trees. Phytobiomes Journal, 5(3), 305–315. 10.1094/PBIOMES-07-20-0049-R

Zandalinas, S. I., Balfagón, D., Arbona, V., Gómez-Cadenas, A., Inupakutika, M. A., & Mittler, R. (2016). ABA is required for the accumulation of APX1 and MBF1c during a combination of water deficit and heat stress. Journal of Experimental Botany, 67(18), 5381–5390. 10.1093/JXB/ERW299

Zhang, Y., Trivedi, P., Xu, J., Caroline Roper, M., & Wang, N. (2021). The Citrus Microbiome: From Structure and Function to Microbiome Engineering and beyond. Phytobiomes Journal, 5(3), 249–262. 10.1094/PBIOMES-11-20-0084-RVW

Zhang, Y., Xu, J., Riera, N., Jin, T., Li, J., & Wang, N. (2017). Huanglongbing impairs the rhizosphere-to-rhizoplane enrichment process of the citrus root-associated microbiome. Microbiome 2017 5:1, 5(1), 97-. 10.1186/S40168-017-0304-4

Zhu, R., Cao, Y., Li, G., Ying, Lianju, G. ·, Ning Bu, M. ·, Hao, L., Zhu, R., Cao, · Y, Li, · G, Guo, · Y, Ma, L, Bu, · N, Hao, · L, Cao, Y., Li, G., Guo, Y., Ma, L., & Bu, N. (2021). Paraburkholderia sp. GD17 improves rice seedling tolerance to salinity. Plant and Soil 2021 467:1, 467(1), 373–389. 10.1007/S11104-021-05108-3

