## Supplementary Figures for "Root-associated microbial community recruitment in two citrus rootstocks subjected to water and salinity stresses"

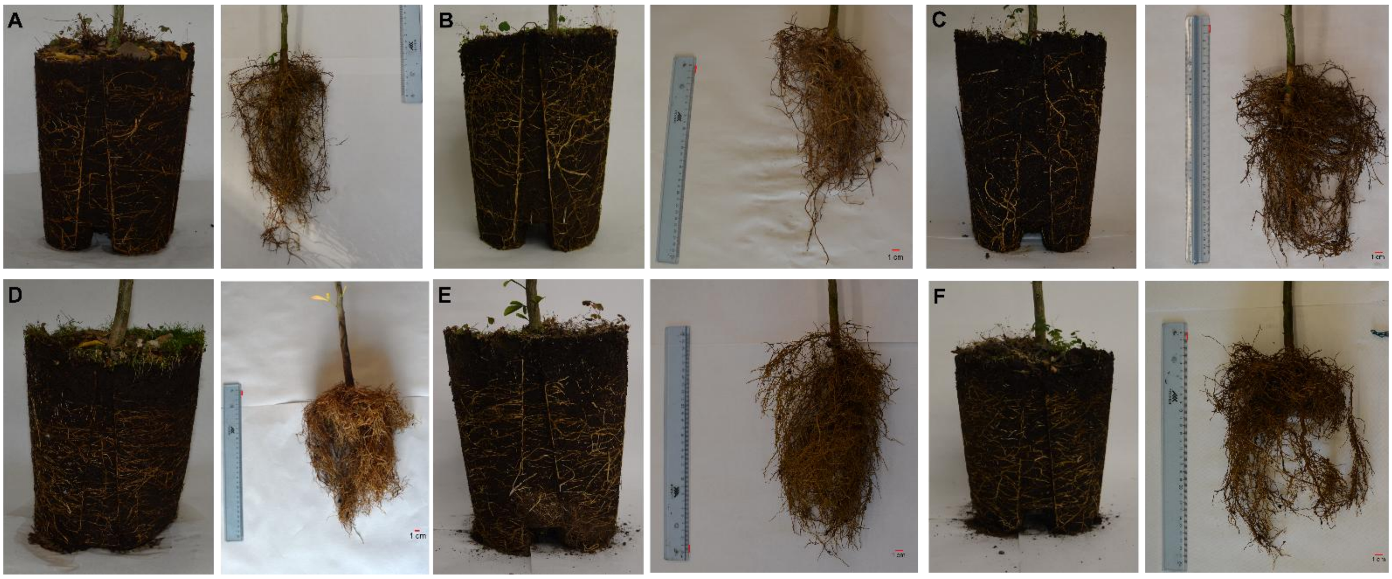


**Supplementary Figure S1.** Representative visualizations of root systems for each genotype and treatment: BIT_CK (A, B), BIT_WS (C, D), BIT_SS (E, F), CRZ_CK (G, H), CRZ_WS (I, L), and CRZ_SS (M, N). Each panel includes two representations, with roots shown embedded in the soil block (left) and after washing (right), representing the corresponding bare root system.

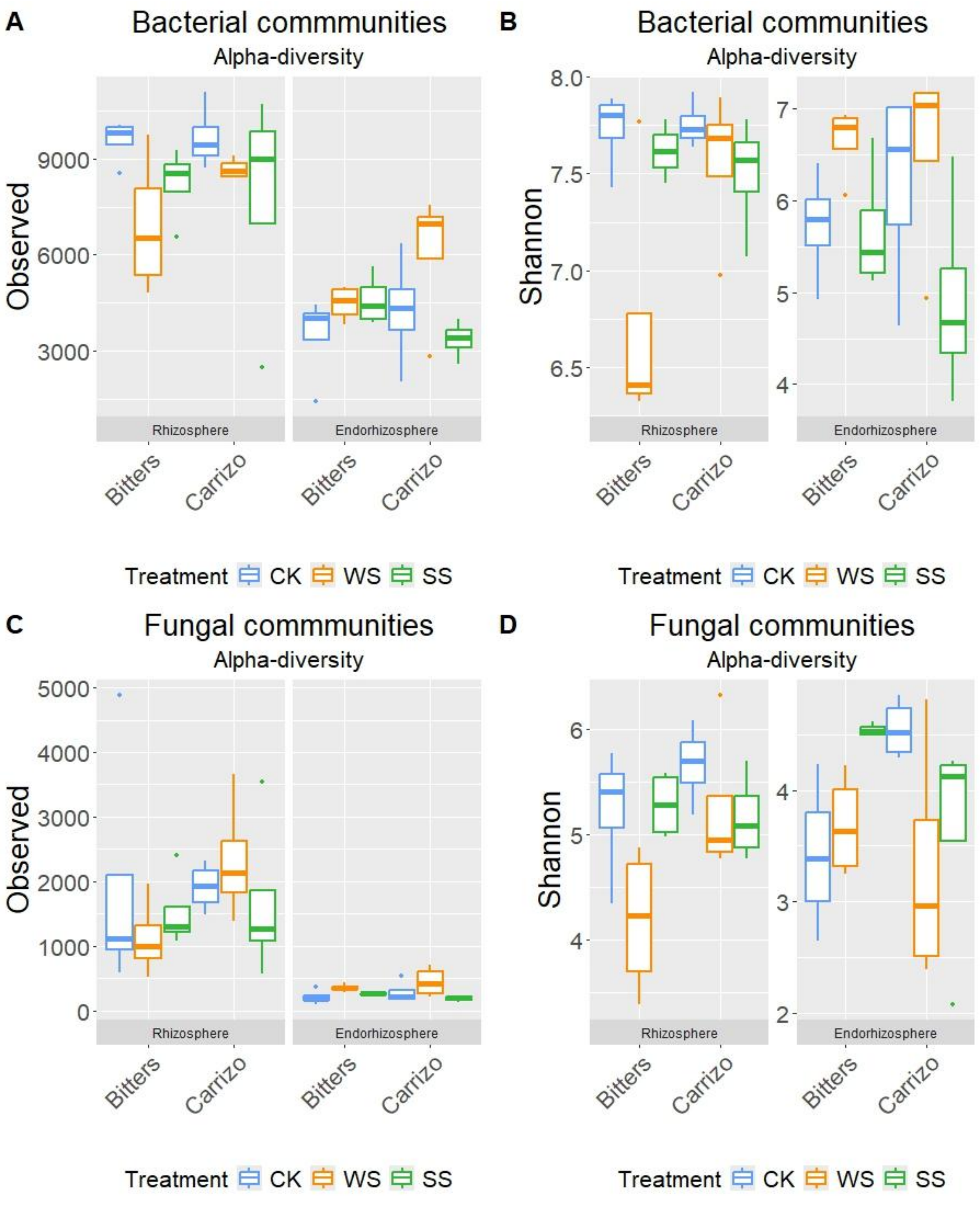


**Supplementary Figure S2.** Alpha diversity estimations of the (A,B) bacterial and (C,D) fungal communities using (A,C) Observed and (B,D) Shannon indices, for each samples (CK, control; WS, water-stressed and SS, salinity-stressed plants) grouped by rootstock (Bitters and Carrizo).

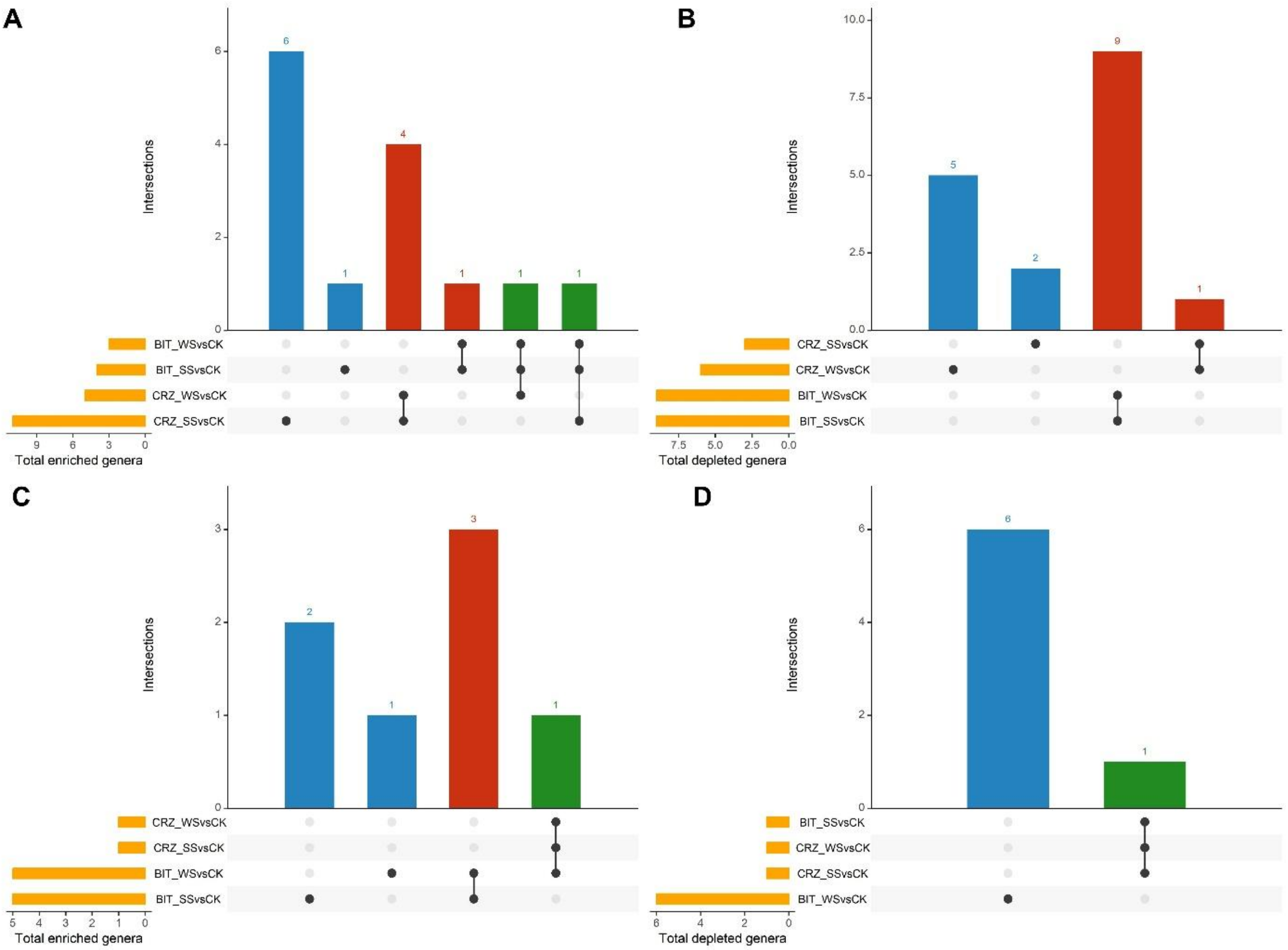


**Supplementary Figure S3.** UpSet plots showing the number of (A, C) enriched and (B, D) depleted fungal genera in each treatment compared with their respective untreated controls (horizontal bars), as well as unique or shared genera across each comparison indicated by individual or connected points, respectively.

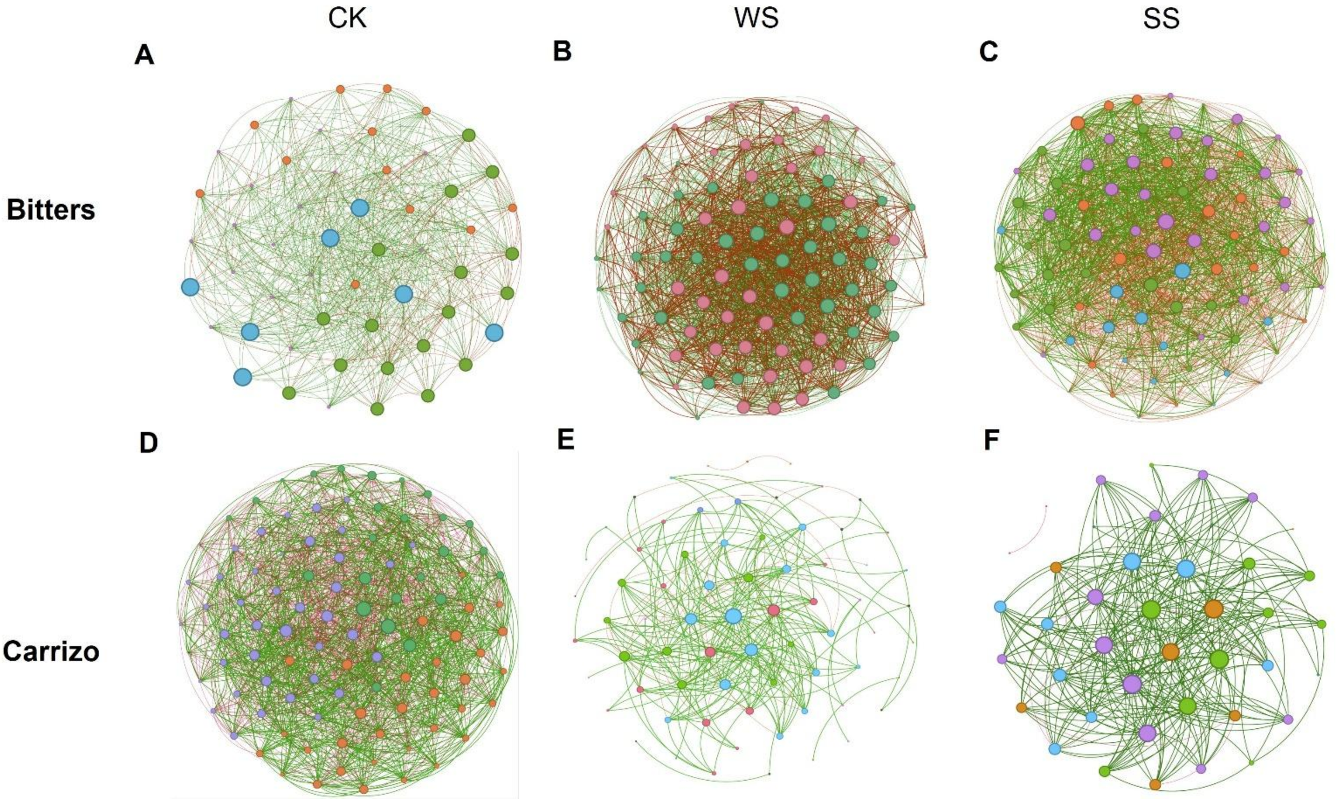


**Supplementary Figure S4.** Co-occurrence networks of fungal communities in control (CK), water-stressed (WS) and salinity-stressed (SS) plants of Bitters (A-C) and Carrizo (D-F) genotypes, respectively
